# Divergent successional patterns and infection dynamics in virion and transcriptionally active soil viral communities following phosphorus amendment and wet-up

**DOI:** 10.64898/2026.04.11.717596

**Authors:** Grant Gogul, G. Michael Allen, Ikaia Leleiwi, Steven J. Blazewicz, Jennifer Pett-Ridge, Joanne B. Emerson, Gareth Trubl

## Abstract

Viruses are key regulators of terrestrial carbon, nitrogen, and phosphorus cycling, yet how environmental perturbations structure viral activity remains poorly resolved. Rewetting of seasonally dry soils triggers rapid microbial and viral responses, but the relationships between virion-associated and transcriptionally active viral communities, and the role of phosphorus in these dynamics, remain unclear. Here, we integrated viromics, metatranscriptomics, environmental DNA (eDNA), and amplicon sequencing to track viral succession and virus–host interactions over three weeks following soil rewetting, with and without phosphorus amendment. We identified 13,840 viral populations (vOTUs), of which 3,803 were transcriptionally active, representing ongoing infections. Wet-up significantly altered virion and transcriptionally active viral communities, while phosphorus selectively influenced prokaryotic and transcriptionally active viral communities but not virion composition. Virus–host linkages were predicted for 32% of vOTUs, with transcriptionally active bacteriophages infecting *Actinomycetota* increasing under phosphorus amendment. Following wet-up, virion abundance decreased ∼3-fold while virocells increased ∼5-fold, indicating a shift from viral persistence in dry soils to active infection. Phosphorus further enhanced virocell abundance. eDNA captured rapid viral turnover and revealed transient dynamics not resolved by viromes or metatranscriptomes alone. Together, these results demonstrate that soil viral communities are structured by distinct but complementary molecular pools that operate over different ecological timescales. Wet-up activates a reservoir of persistent virions, while phosphorus availability regulates infection dynamics and host–virus coupling. These findings highlight viruses as dynamic drivers of microbial turnover and nutrient cycling following environmental perturbation, advancing a more predictive understanding of soil ecosystem responses to changing resource availability.

## Introduction

Soils harbor a diverse array of viruses, and those that infect microbial hosts are presumed to play key roles in terrestrial biogeochemical cycles, such as the carbon and phosphorus cycles ^1–5^. Soil viruses can rival microbial host abundances, with up to 10^10^ viral particles (virions) per gram of soil ^6–9^, highlighting their potential to exert top-down control on microbial host populations via mortality influencing microbial community structure, nutrient availability, and soil organic matter quality ^10–12^. The structured and heterogeneous nature of soil can limit viral dispersion, leading to highly divergent viral communities over time and space ^13–16^. These pronounced spatiotemporal trends have made it challenging to determine which viruses are actively infecting hosts in soil and to understand how these infections shape virus–host interactions and terrestrial biogeochemical cycling.

An additional layer of complexity arises from the presence of environmental DNA (eDNA; also known as relic or extracellular DNA), which includes DNA originating from live, dead or lysed organisms and can represent a large fraction of total nucleic acids recovered from soils ^15,17,18^. eDNA pools are governed by the balance between biological inputs and degradation processes and can persist independently of living or transcriptionally active communities ^19^. eDNA pools may not bias diversity estimates; however, they can influence the interpretation of community composition and dynamics ^19,20^. As eDNA can substantially alter inferred diversity and taxon-specific abundance patterns in some circumstances, eDNA represents a distinct and ecologically relevant nucleic acid pool that should be considered when interpreting soil microbial and viral datasets ^20^.

Mediterranean grasslands provide an ideal system for studying viral community responses to precipitation. Pronounced seasonal shifts in rainfall, from hot, dry summers to cool, wet winters, generate large natural variation in soil water potential over the course of the year. The dry season reduces microbial activity and viral detection, while initial wet season precipitation stimulates microbial and viral activity, driving community succession ^15,16,21–25^. Following the initial wet-up, DNA viral communities turn over more rapidly than their microbial hosts ^15^, contributing to a surge in microbial mortality within 24 hours of rewetting ^22,25^ and shifts in DNA ^15,25^ and RNA ^26^ viral community composition and richness. However, these patterns for DNA viruses have only been documented over relatively short timescales (≤10 days), leaving longer-term impacts on viral communities unresolved.

Phosphorus is a fundamental element for life and, in many soils, is a limiting nutrient ^27^, influencing microbial growth rates, ecological interactions, and virus–host dynamics ^28,29^. Inorganic forms of phosphorus in soil, such as phosphate, can be incorporated into phospholipids, ATP, and nucleic acids by soil microorganisms. However, because virions are mainly composed of nucleic acids, they exhibit a lower carbon-to-nitrogen-to-phosphorus (C:N:P) stoichiometric ratio than their microbial hosts ^30,31^. Virions’ relatively high phosphorus demand has led to the hypothesis that viruses may immobilize and deplete microbial necromass of phosphorus ^30^. In culture, phosphorus availability influences how viruses manipulate their host metabolism by increasing nitrogen assimilation, fatty acid degradation, and resource acquisition ^32^, while reducing production of proteins involved in *de novo* nucleotide synthesis ^33^. Viruses rely on host phosphorus pools for replication; therefore, availability can alter viral infection rates and virion production, ultimately leading to shifts in microbial host populations. Previous studies have shown that phosphorus-limited environments lead to a lower burst size ^34^ and decrease viral transcription during infection ^32^, yet the effects of phosphorus availability on soil DNA viral communities and virus infection dynamics remain unreported.

Here, we sequenced temporally resolved viromes, metatranscriptomes, 16S rRNA gene amplicons, and eDNA (< 0.02 µm size fraction) from California grassland soil microcosms over three weeks to investigate virus–microbe interactions following a simulated precipitation event and a phosphorus amendment. This integrated approach allowed us to track both virions (via viromics) and transcriptionally active viral infections (via metatranscriptomics) over time, revealing how viral community and virus–host dynamics shift in response to environmental perturbation. Here, we address three key gaps: (1) how virion-associated and transcriptionally active viral communities differ, (2) how these communities respond over ecologically relevant timescales following rewetting, and (3) how nutrient availability, specifically phosphorus, regulates viral infection dynamics.

## Materials and methods

### Soil collection and experimental setup

Soil was collected prior to the first seasonal rainfall in Autumn on October 3, 2020, from the Buck pasture plot at the Hopland Research and Extension Center (HREC; Mendocino County, CA; 39.001767° N, 123.069733° W) at an elevation of 1,066 feet and a depth of 0–10 cm. Soil was sieved (2.0 mm), homogenized, and transported to Lawrence Livermore National Laboratory.

Microcosms were established by adding 206 g (± 0.5 g) of soil to 1.9 L acid-washed and autoclaved jars and received one of two treatments: (1) deionized water (dH_2_O; pH 5.5) ^23^; or (2) dH_2_O + KH_2_PO (50 µM; 1.37 μg P g^-1^soil). Water was evenly applied to the soil using a syringe to achieve 30% GWC. Treatments were replicated six times for each of the three time points (days 7, 14, and 21 post wet-up), totaling 36 microcosms. Microcosms were sealed and incubated at ∼23°C.

Initial dry soils and microcosms were destructively sampled (39 total samples) on days 7 (T1), 14 (T2), and 21 (T3) post wet-up. Soils were homogenized and subsampled for physiochemical measurements and nucleic acid extractions. Detailed procedures are provided in Supplementary Methods.

### Soil virome processing and DNA extraction

Viral like particle (VLP) were extracted from10 g of soil thawed at 4°C for 1 hour following previously established protocols with minor amendments ^35,36^. Briefly, soil was washed with AKC’ buffer, shaken, vortexed, and centrifuged. The resulting supernatant was filtered through a 0.22 µm filter to remove most cellular debris and concentrated using a pretreated 100 kDa Amicon Ultra-15 filter. The Amicon filter was further washed three times with AKC’ buffer. The VLP concentrate was DNase treated to remove DNA not encapsidated by a VLP. VLPs were then precipitated using FeCl_3_ and DNA extracted using a phenol-chloroform-based method and quantified (Supplementary Table 2) ^37,38^.

Extracted DNA was shipped to the Joint Genome Institute (JGI; Berkeley, CA) for library preparation and Illumina sequencing. Initially, DNA libraries were prepared for 16 of the 39 samples using the Accel-NGS 1S kit (Swift Biosciences; Supplementary Table 1) and sequenced using the Illumina NovaSeq platform (2×151bp), generating ∼300 million paired-end reads per sample. The Accel-NGS 1S kit was chosen due to its ability to amplify both single-stranded and double-stranded DNA ^36,39,40^. Due to availability issues of the Accel-NGS 1S kit, the SRSLY NGS library Prep Kit (Claret Biosciences, Santa Cruz, CA, USA) was used for 38 of the 39 samples. One sample (T3H218O_A) was not used due to insufficient DNA. In total, 54 viromes were generated (see Supplementary Table 1). See Supplementary Methods for detailed VLP concentration, DNA extraction, and library preparation protocols.

### Virome read processing and vOTU identification

All sequencing reads (virome, metatranscriptome, and eDNA) were quality filtered using BBDuk ^41^, assembled using SPAdes single cell mode (v. 3.15.4) ^42,43^, mapped to reference sequences using Bowtie2 (v. 2.5.2) ^44^, with alignments processed with SAMtools (v. 1.19) ^45,46^, final abundance tables of mapped reads were generated using CoverM (v. 0.6.1) ^47^.

For samples prepared with the SRSLY library kit, reads were additionally trimmed according to manufacturer recommendations.

Viral contigs were identified using VirSorter2 (v. 2.2.4) ^48^ and geNomad (v. 1.7.1) ^49^, assessed with CheckV (v.1.0.1) ^50^, and retained following previously established criteria ^48^. Briefly, viral contigs were filtered based on presence or absence of viral and host genes, viral score, and length of the viral contig. To capture both ssDNA and dsDNA viruses, contigs were filtered by length, retaining contigs >10 kbp and 1–10 kbp, with the latter restricted to those classified as *Monodnaviria* by geNomad. Resulting filtered viral contigs were clustered using established parameters (>95% ANI, over 85% of the contig) to generate a dereplicated set of viral populations (vOTUs) ^51^. vOTUs were additionally screened for auxiliary metabolic genes (AMGs) using DRAM-v. Detailed protocols, parameters, and filtering criteria are provided in the Supplementary Methods.

### Bulk soil RNA extraction and read processing

Total RNA was extracted from soil using a phenol–chloroform-based protocol from previously published methods ^52^. RNA was cleaned up using the Qiagen All-Prep kit (Qiagen). RNA library preparation and sequencing methods were previously reported ^26^, including DNase treatments, rRNA depletion, and library preparation using NEBNext Ultra II RNA Library Prep Kit (NEB, Ipswich, MA, USA). Libraries were sequenced on an Illumina HiSeq 2500 platform (a 2×150 bp), generating ∼350 M paired-end reads per sample.

Reads were quality filtered and trimmed to remove adapters, barcodes, primers, and PhiX contamination, followed by quality trimming. Detailed protocols and filtering criteria are provided in the Supplementary Methods.

### vOTU relative abundances in viromes and metatranscriptomes

Quality-filtered virome reads from the SRSLY library preparation kit were mapped to vOTUs, and abundance tables were generated using previously established thresholds of >90% read identity covering at least 75% of the contig ^5,39^ and normalized by contig length and sequencing depth, yielding normalized abundance tables for downstream analyses (Supplementary Table 4).

Quality-processed metatranscriptomic reads were mapped to DNA vOTUs and predicted genes using Prodigal-gv (v.2.11.0-gv)^49,53^ to identify transcriptionally active vOTUs ^3,54^. Coverage and counts tables for mapped metatranscriptomic reads were generated requiring a minimum of 90% read identity ^54^. For each sample, if the total number of reads mapped to a putative gene across triplicate metatranscriptomes were fewer than four, the abundance of that gene was set to zero ^54^. vOTUs were considered transcriptionally active if at least one gene was expressed per 10 kbp of the vOTU, otherwise the relative abundance of the vOTU was set to 0 ^3,54^. Transcriptionally active vOTUs were normalized by vOTU length and library size (Gbp), generating the abundance table for downstream analyses (Supplementary Table 5). Detailed mapping parameters, thresholds, and filtering criteria are provided in the Supplementary Methods.

### Environmental DNA extraction, sequencing, and analysis

eDNA was extracted from 5 g of soil thawed at 4 °C for 1 hour. Soil was washed twice with 10 mM Tris-HCl (pH 8.0) by shaking and alternating between vortexing and hand mixing. Samples were then centrifuged and sequentially filtered through 0.22 µm and 0.02 µm filters to remove cellular and viral particles. eDNA was precipitated using isopropanol as described above. eDNA concentrations were quantified and used for library preparation with the NEBNext Ultra DNA Library Preparation Kit (New England Biolabs, Ipswich, MA, USA) sequenced on an Illumina HiSeq platform (2×150 bp).

Raw eDNA reads were quality filtered and mapped as previously described and vOTU abundance tables of mapped reads were generated requiring a minimum of 90% read identity. vOTUs were considered detected in a sample only if the summed number of reads mapped across triplicate samples exceeded three reads using the counts table generated from CoverM. The mean coverage tables were the filtered for vOTUs detected in each sample and normalized by total gigabases (Gbp) sequenced per eDNA library resulting in an abundance table representing the vOTU coverage normalized by contig length and sequencing depth (Supplementary Table 3). Detailed protocols, parameters, and filtering criteria are provided in the Supplementary Methods.

### Host prediction

Virus hosts were predicted using iPHoP (v1.3.3) ^55^ with the iPHoP_db_Aug23_rw database (–min_score 75). Genus-level assignments were retained only when the confidence score was ≥ 90%, otherwise predictions at the family or higher taxonomic level were used ^40^. Host-associated viral community composition was summarized by aggregating vOTU relative abundance at the host phylum level for viromic and metatranscriptomic datasets. For visualization dominant host phyla were retained and less abundant groups were grouped into an “Other” category. Detailed filtering, normalization, and plotting procedures are provided in the Supplementary Methods.

### Amplicon processing

DNA was extracted from 0.5 g of soil using the Powersoil Pro kit (Qiagen, Hilden, Germany). The V4 region of the 16S rRNA gene was amplified using primers 515F–Y and 806R ^56,57^, with library preparation and PCR conditions as previously described by Sieradzki *et al.* ^26^. Amplicons libraries were sequenced on an Illumina MiSeq platform (2×250 bp) with ∼25% phiX spike-in.

Raw reads were quality filtered, adapter-trimmed, and processed. DADA2 (v.1.30.0) ^58^ was used to infer amplicon sequence variants (ASVs). Taxonomy was assigned using the GTDB_bac120_arc53_ssu_r214_genus reference database, and ASVs lacking a kingdom-level assignment were removed.

Samples were rarefied to the 10th percentile of sequencing depth. Samples with insufficient sequencing depth (T1H218O_A, T1H2O_A, T1H2OP_C) were excluded from downstream analyses (Supplementary Table 9). Detailed processing parameters are provided in the Supplementary methods.

### Virion and virocell abundance predictions

Virion and virocell abundances were estimated following approaches adapted from prior work ^25^. Briefly, virion abundance was inferred by scaling the proportion of virome reads mapping to vOTUs by the total DNA extracted to estimate viral DNA (equation 1), which was converted to particles using average vOTU genome weights (equation 2).

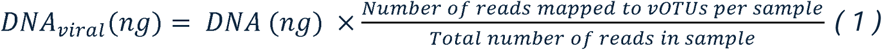

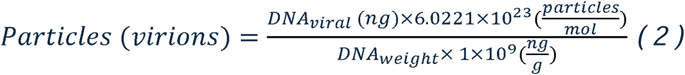

Virocell abundance was estimated by scaling the proportion of metatranscriptomic reads mapping to DNA vOTUs by the total RNA extracted per sample to estimate viral RNA (equation 3), which was converted to viral particle equivalents using average vOTU weights (equation 4). At the viral community level, the number of viruses produced per infected cell (burst size) can vary, so a range of viruses per cell was used (1–200 viruses per cell; equation 5).

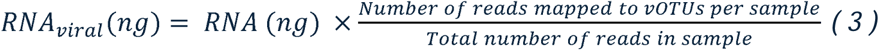

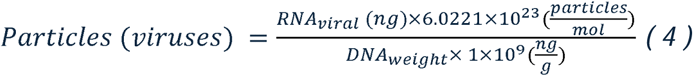

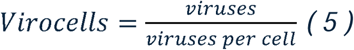

All estimates were normalized to dry soil mass to enable comparisons across samples. For detail explanations of calculations see Supplementary Methods.

### Statistical analyses

All statistical analyses were conducted in R (v.4.4.2) ^59^ and rstatix (v.0.7.2) ^60^. Viral community beta-diversity was assessed using Bray-Curtis dissimilarity on Hellinger transformed abundance tables and tested using permutational multivariate analysis of variance (PERMANOVA), with pairwise PERMANOVA comparisons when appropriate. Host–virus interaction analyses were evaluated using distance-based redundancy analysis (dbRDA), with the significance assessed by permutation-based analysis of variance (ANOVA).

Difference in virion and virocell abundances across time points and phosphorus treatments were assessed using linear models and ANOVA with post hoc Tukey’s HSD tests. To determine if virocell predictions exhibited similar temporal trends for different burst sizes, significance was determined using generalized additive model (GAM). Nucleic acid concentrations were normalized to the maximum mean value across time points, and temporal changes were assessed using linear models with parametric or non-parametric tests as appropriate.

A heatmap of eDNA abundances was generated from log10-transformed normalized abundance. All figures were generated using ggplot2 (v.3.5.1) ^61^, ggpubr (v.0.6.0) ^60^, and patchwork (v.1.3.0) ^62^. Detailed statistical procedures, model specifications, and software parameters are provided in the Supplementary Methods. Results of statistical analyses are further documented in Supplementary Tables 10–15.

## Results and discussion

### Virion and transcriptionally active viral communities were compositionally distinct

To investigate the response of DNA viruses to soil rewetting and phosphorus amendment, we leveraged viromics and metatranscriptomics in a replicated time-series microcosm study. We identified 13,840 viral operational taxonomic units (vOTUs) in the virome, the majority were dsDNA viruses (13,617), with 223 ssDNA viruses. Using metatranscriptomics, we identified 3,803 transcriptionally active vOTUs (27.5% of total vOTUs) based on viral gene detection, 38 were ssDNA viruses. Transcriptionally active vOTUs were defined based on detection of viral gene expression across the dataset. The overall viral community composition was significantly structured by its source (virome vs. metatranscriptome) as determined by PERMANOVA (*p* < 0.05, *R^2^* = 0.035; Fig.1A). Most vOTUs (72.5%) were unique to the virome (i.e., not detected as transcriptionally active), with a three-fold higher viral richness compared to transcriptionally active viruses (Fig. 1B and C; Supplementary Fig. 1A). These findings align with a previous report showing that approximately 22% of vOTUs were identified as newly replicated virions using ^18^O-stable isotope probing following soil rewetting ^63^. To confirm these results were not driven by sequencing depth (i.e., the lower number of vOTUs detected in metatranscriptomes was not due to sequencing effort), we generated accumulation curves that demonstrated our sampling and sequencing depth sufficiently captured the viral diversity in both the virome and transcriptionally active viral communities (Fig. 1D).

**Figure 1:**
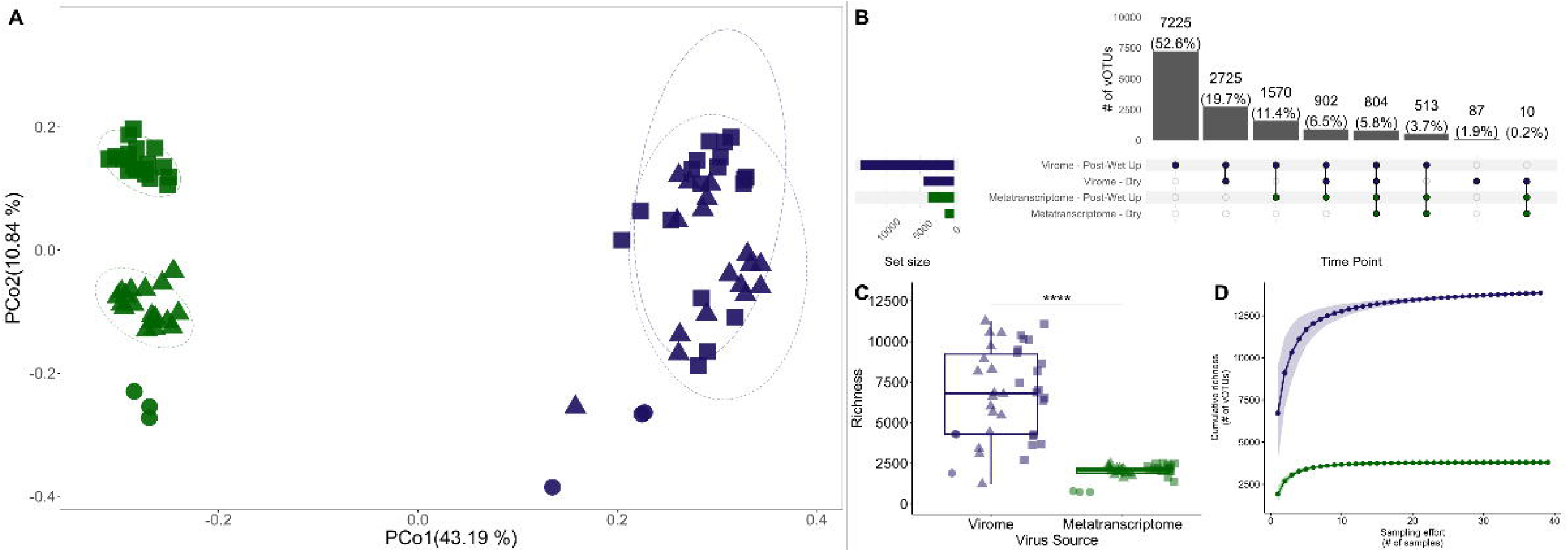
Virion-associated and transcriptionally active viral communities represent distinct ecological pools. (A) Principal Coordinates Analysis of Bray-Curtis dissimilarities, colored by viral community type with shapes representing treatments. (B) An UpSet plot illustrating detection patterns of vOTUs present in at least one replicate. Vertical bar plots indicate the number of shared vOTUs across conditions indicated by highlighted circles and set sizes (horizontal bars) indicate the total number of unique vOTUs per condition. (C) Boxplot showing viral richness for both the virome and transcriptionally active viral community. The horizontal line within each box indicates the median richness and each point represents the vOTU richness of an individual sample. (D) Accumulation curve showing cumulative viral richness per number of samples for the virome (purple) and transcriptionally active viruses (green). * *p* < 0.05, ** *p* < 0.01, *** *p* < 0.001, **** *p* < 0.0001 (Dunn’s test).

The significant divergence between transcriptionally active and virion-associated viral community indicates that only a subset of soil vOTUs exhibit detectable viral gene expression within host cells at any given time. These results demonstrate that virion-associated and transcriptionally active viral communities represent fundamentally different ecological pools, capturing viral persistence versus active infection. Transcriptionally active vOTUs were stringently defined such that only vOTUs with sufficient gene-level expression across the dataset were considered. As a result, metatranscriptomes reflect a conservative subset of viral populations undergoing detectable intracellular transcription near the time of sampling, rather than the full diversity of viruses present in the soil. In contrast, virome-derived vOTUs represent intact virions present in the soil, which can remain detectable for longer periods than viral transcripts and thus reflect vOTUs present over a broader temporal window. This difference in detection reflects both biological processes and methodological resolution, as viral mRNA is short-lived and requires ongoing transcription to be captured ^64,65^, whereas virions can persist in soils for days to weeks following production ^66^. This longer environmental residence time of virions has been proposed to result in partial carryover of virion-associated vOTUs across time, such that viromes may reflect vOTUs produced during prior infection events as well as those generated contemporaneously with sampling ^15,63,67^. Similar time-integrative effects have been described for other biological pools in soils, including microbial communities, where slowly turning-over components can obscure short-term dynamics ^68^.

Most vOTUs detected in viromes after wet-up were not present in dry soils, indicating substantial viral turnover over the span of this experiment, consistent with new virion production following rewetting, even though many of these vOTUs were not detected in the metatranscriptomes. The decoupling of virion detection and transcriptional activity is expected given the amount of soil used, time between sampling for each method, and conservative thresholds applied to identify transcriptionally active vOTUs. Additionally, viral transcription occurring in a limited fraction of infected cells, at low expression levels, or outside the immediate sampling window may therefore fall below detection thresholds in metatranscriptomes. Although virions in soils are not permanently stable, their environmental residence time can nonetheless dampen or obscure short-term changes in viral activity, similar to patterns observed in microbial communities ^17,20^. Distinguishing between virion-associated and transcriptionally active viral fractions using complementary approaches is essential for resolving soil viral dynamics, as their integration provides a more accurate view of viral turnover and the subset of viral populations most directly influencing microbial hosts and biogeochemical processes over time and space, despite inherent limitations of metatranscriptomes.

### Virion and transcriptionally active viral communities exhibited divergent temporal responses to soil wet-up

To capture longer-term viral community dynamics following soil rewetting, we extended upon previous studies of up to 10 days ^15,25,63^ by monitoring both the virome and transcriptionally active viral communities over a three-week period. Both communities displayed strong responses to wet-up, which served as a significant driver of community structure (PERMANOVA, *p* = 0.001, *R^2^* = 0.219 for viromes; *p* = 0.001, *R^2^*= 0.167 for transcriptionally active viruses; Fig.2A, B). Notably, transcriptionally active viral richness increased after the first week, similar to prior observations in the virome ^15^, although virome richness here did not increase until the second week (*p <* 0.01; Fig. 2C, D). These responses are consistent with the expectation that rapid infection and replication dynamics occur immediately after wet-up ^69^.

**Figure 2:**
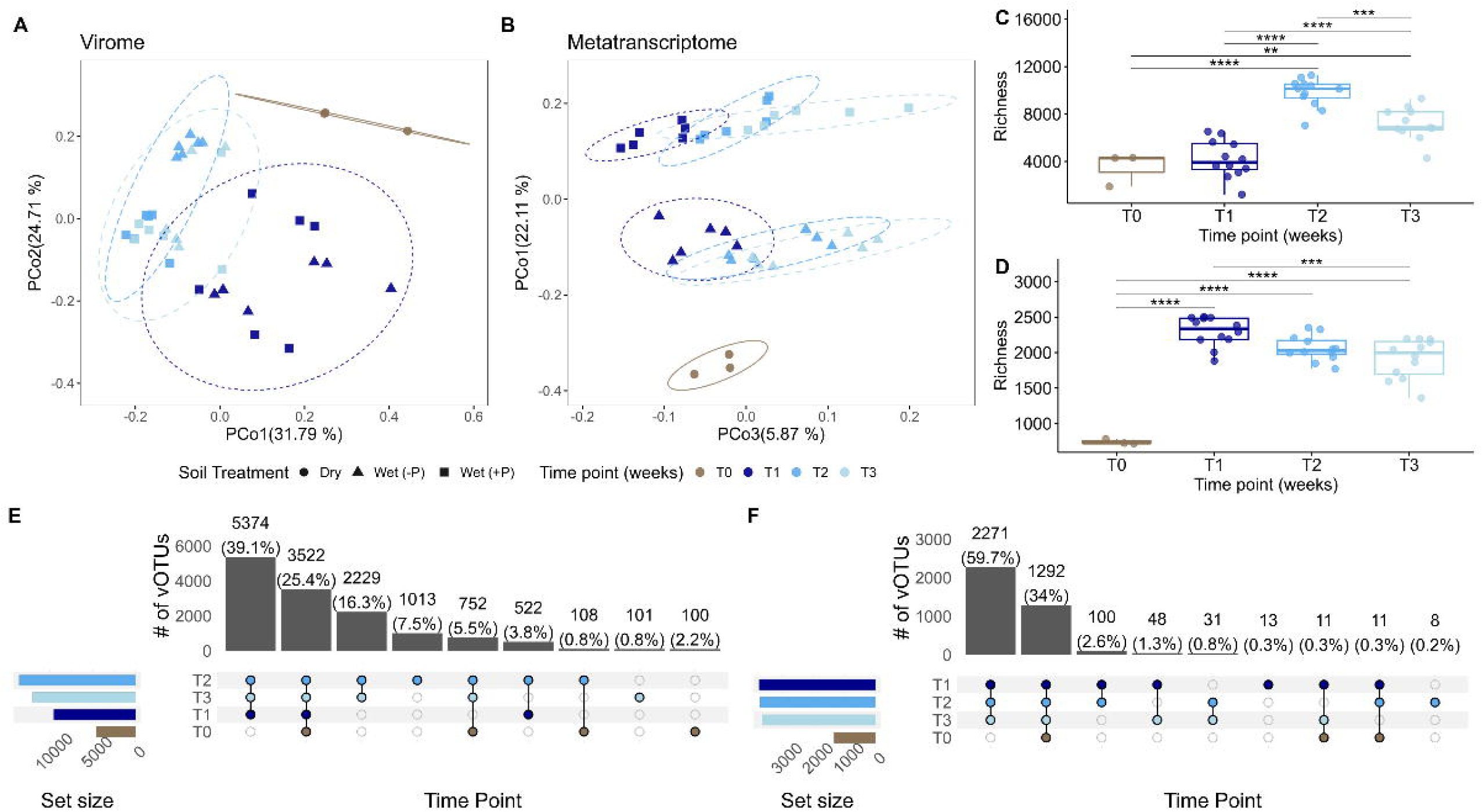
Divergent temporal dynamics of virion and active viral communities following wet-up. Principal Coordinates Analysis of Bray-Curtis dissimilarities of the (A) virome and (B) transcriptionally active viral community; colors and dashed lines indicate sampling time points, and shapes represent soil type. Ellipses represent 95% confidence region around each time point. Boxplots show viral richness over time in (C) viromes and (D) transcriptionally active viral communities, with colors representing time point; horizontal lines within each box are the median viral richness and each point is the viral richness of an individual sample. UpSet plots illustrate vOTU detection patterns over time for the (E) viromes and (F) transcriptionally active viral communities; colors represent time points, vertical bar plots are the number of shared vOTUs in at least one replicate among time points indicated by filled circles and set sizes (horizontal bars) show the number of unique vOTUs detected per time point. The UpSet for the virome and transcriptionally active viruses are only showing groups with greater than 85 or 8 shared vOTUs, respectively. T0 = dry soil, T1 = 1 week, T2 = 2 weeks, T3 = 3 weeks post–wet-up. * *p* < 0.05, ** *p* < 0.01, *** *p* < 0.001, **** *p* < 0.0001 (ANOVA and Tukey’s HSD).

To assess viral community dynamics throughout the three-week experiment, we examined community composition across post-wet-up samples. A large portion of vOTUs remained detectable across all three weeks for both the virome (∼39%) and the transcriptionally active viral community (∼60%), indicating a stable community of viruses (Fig. 2E, F). Despite the temporal overlap of vOTUs, the beta-diversity patterns revealed ongoing community succession patterns initially triggered by rewetting then stabilizing by the second week, with no significant difference between the second and third weeks (pairwiseAdonis, Supplementary table 10 & 11). Consistent with a parallel study of RNA viruses from the same dataset, wet-up was a main driver of viral community structure ^26^, but RNA viral communities remained relatively stable for the three weeks following wet-up, unlike the DNA viral community composition here that continued to shift through the second week (Supplementary table 10). This pattern may arise from differences in host associations, as RNA viral dynamics may be driven by mycoviruses ^26^, whose fungal host remain relatively stable following wet-up ^52^, whereas the DNA viral community primarily infect bacteria which exhibit taxon-specific responses to soil rewetting ^22,52^. Our results demonstrate soil rewetting triggered rapid viral community succession followed by stabilization. The duration and stability of these patterns under longer periods (beyond three weeks) or subsequent drying remain open questions for future studies. This pattern is consistent with a rapid activation phase followed by stabilization, indicating that viral dynamics operate on short ecological timescales following environmental perturbation.

### Phosphorus amendment shifted prokaryotic and transcriptionally active viral community composition and enhanced virocell production

Phosphorus availability significantly shaped viral activity and microbial communities following soil rewetting. While phosphorus did not significantly affect the virome viral community structure (Fig. 3A), there was a significant difference within the transcriptionally active virus and microbial (16S rRNA gene) communities, as indicated by PERMANOVA (Fig. 3B, C). Despite these compositional changes, richness did not differ significantly (*p* value > 0.05) between phosphorus amended soils and unamended soils for viromes, transcriptionally active viruses, or prokaryotic communities (Fig. 3D–F). A majority of transcriptionally active vOTUs (96%) were shared between phosphorus-amended and unamended soils (Supplementary Fig. 1), indicating that, despite a significant response to phosphorus, a reservoir of transcriptionally active viral populations was present regardless of phosphorus amendment. Therefore, compositional differences between treatments were driven by differences in transcriptional activity of shared vOTUs (e.g., higher expression under phosphorus amendment relative to unamended conditions), rather than the presence of unique vOTUs. Thus, phosphorus availability altered viral transcription, resulting in significant differences in transcriptionally active community composition.

**Figure 3:**
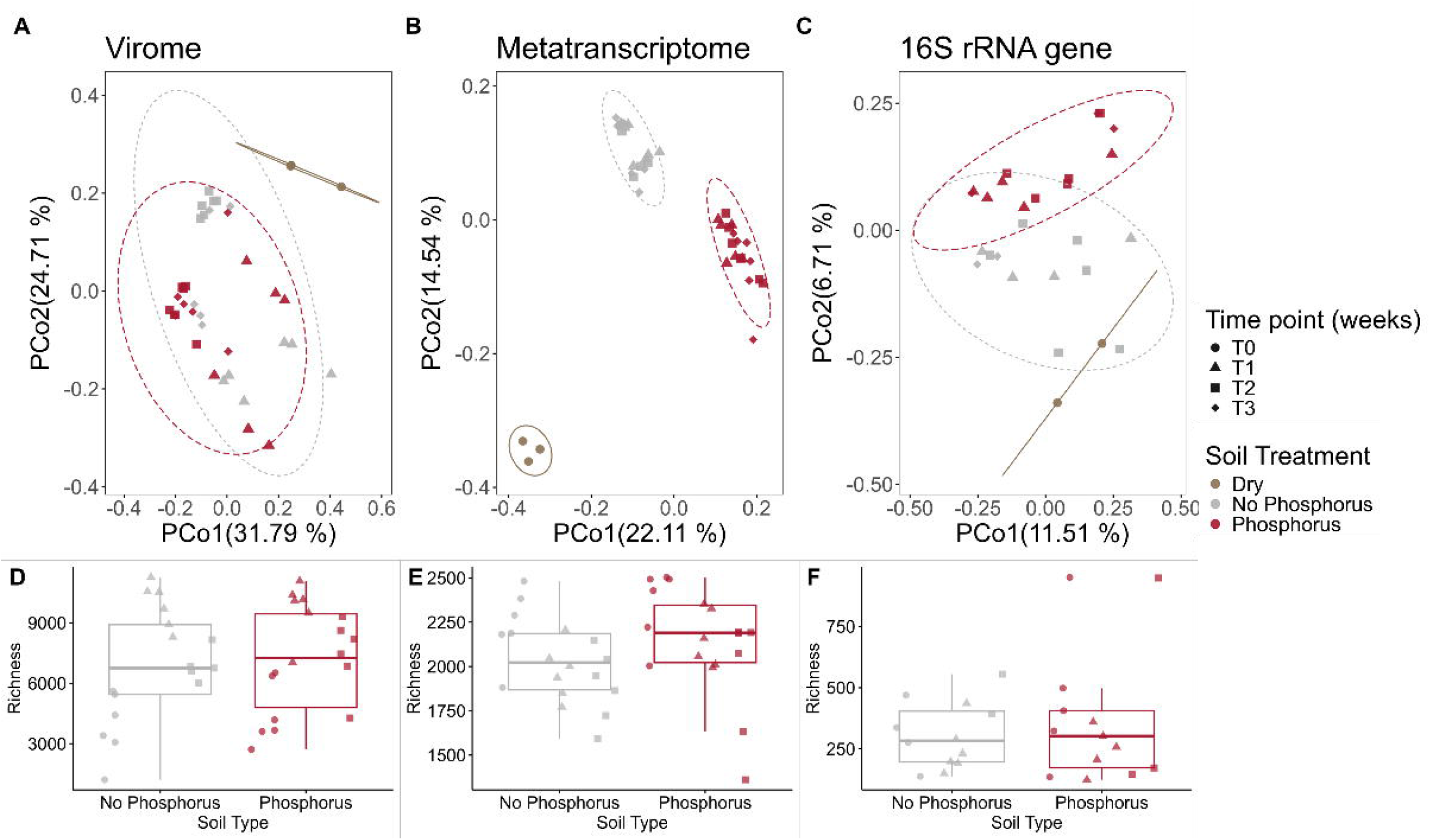
Phosphorus regulates transcriptionally active viral communities but not virions. Principal Coordinates Analyses of Bray-Curtis dissimilarities in (A) virome, (B) transcriptionally active viral, and (C) microbial communities (16S rRNA gene). Colors and dashed lines represent soil type; shapes represent time points. Boxplots show viral richness in (D) viromes, (E) transcriptionally active viral communities, and (F) microbial communities (16S rRNA gene), and horizontal lines indicate medians. Colors denote soil type; each point is an individual sample. T0 = dry soils, T1 = 1 week, T2 = 2 weeks, T3 = 3 weeks post–wet-up. * *p* < 0.05, ** *p* < 0.01, *** *p* < 0.001, **** *p* < 0.0001 (ANOVA and Tukey’s HSD).

To assess the effects of wet-up and phosphorus on viral infection dynamics, we integrated viromic and metatranscriptomic data to estimate the abundances of virions and virocells. Following wet-up, the estimated average number of virions decreased nearly two-fold, while the number of virocells increased approximately six-fold (Fig. 4A, B). These reciprocal trends suggest that virions were either degraded ^66^, assimilated by other organisms ^70,71^, or transitioned into active infections, coinciding with elevated viral transcriptional activity (Fig. 4B). Although phosphorus did not significantly alter estimated virion abundance (ANOVA, *p* > 0.05), it significantly increased the estimated number of virocells across all post–wet-up time points (ANOVA; Tukey’s HSD, *p* < 0.01; Fig. 4A, B), suggesting that phosphorus availability promotes viral infection and transcription. These findings are consistent with prior work showing phosphorus influences viral infection dynamics, including increased RNA bacteriophage (phage) abundance following wet-up ^26^ and enhanced virocell production and transcriptional activity in culture ^32^. Together, these results indicate that viral infection is influenced by phosphorus availability, and that when phosphorus is not limiting, viruses can proliferate and increase the number of infected cells. Supporting the idea that viruses contribute to nutrient turnover and biogeochemical cycling in terrestrial ecosystems through host infection. These results indicate that phosphorus regulates viral infection dynamics primarily through effects on host–virus interactions rather than direct control of virion production.

**Figure 4:**
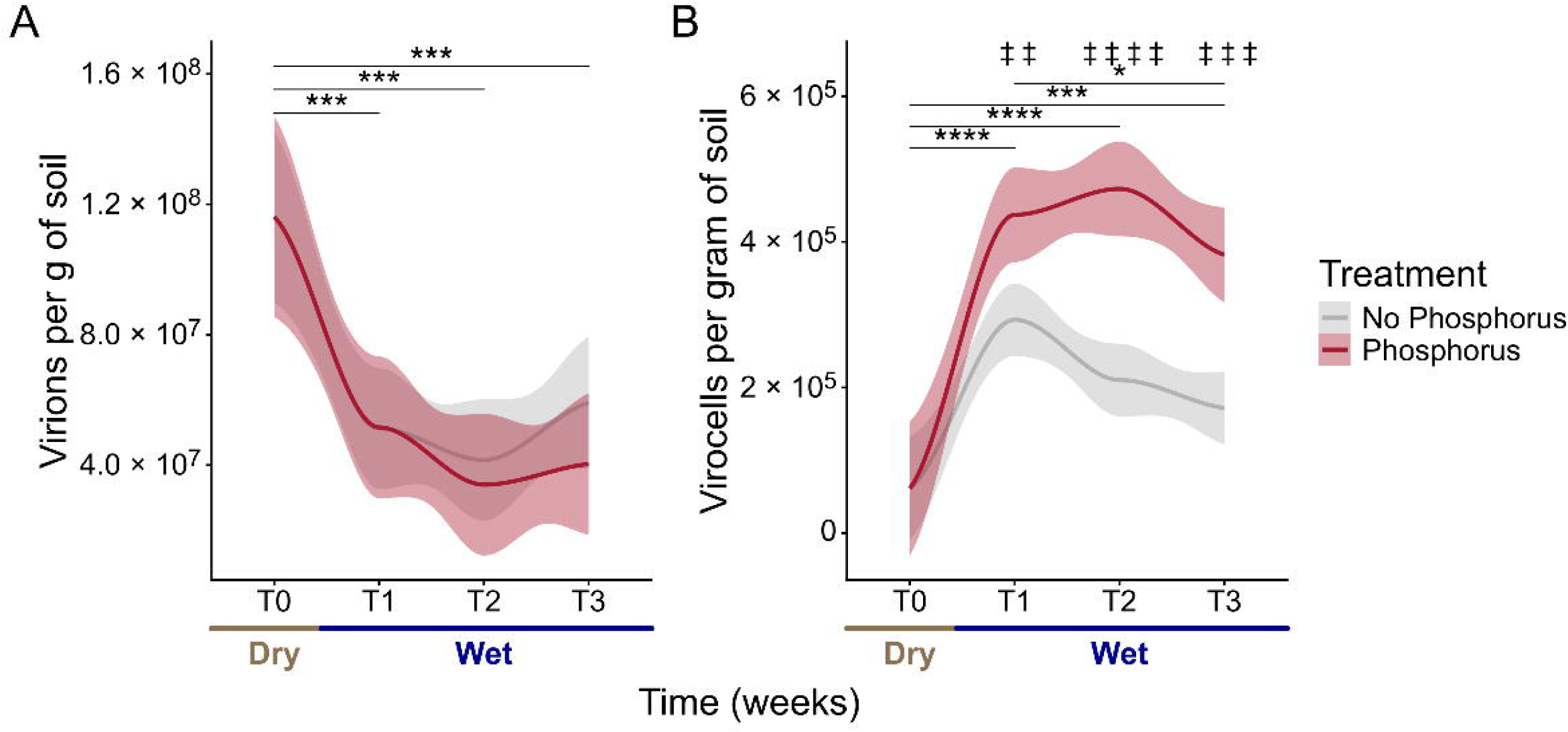
Wet-up shifts viral communities from persistence to active infection, amplified by phosphorus. Ribbon plots show the predicted numbers of (A) virions and (B) virocells per gram of soil, assuming a burst size of 200 virions produced per cell per successful infection. Colored lines represent mean values within each treatment across time; ribbons indicate 95% confidence intervals. Brown line below x-axis indicates dry soil samples, blue line indicates all samples following wet-up. Asterisks denote a significant difference between the average abundance at each time point, while the double daggers indicate significant treatment effects within a time point; T0 = dry soils, T1 = 1 week, T2 = 2 weeks, T3 = 3 weeks post–wet-up. * *p* < 0.05, ** *p* < 0.01, *** *p* < 0.001, **** *p* < 0.0001 (ANOVA and Tukey’s HSD).

While our approach provides insight into viral abundance and activity, these estimates are constrained by methodological limitations and underlying assumptions. Although there is precedent for using viromic DNA yield as a proxy for viral particle abundances ^15^, calculations based on nucleic acid concentrations and read-mapping data have inherent uncertainty. For example, they assume similar DNA extraction efficiency across samples. Moreover, the assumed burst size of 200 used in the virocell estimates was an approximation, as there is no consensus for an average burst size for soil viruses, and burst size likely correlates with environmental conditions ^72–74^. Nonetheless, our estimates of virion abundance per gram of soil were consistent with previously reported values in other soils (2.0×10^7^– 1.0×10^10^ per gram of soil) ^7^, lending confidence to the relative trends observed here. While absolute values should be viewed as approximations until confirmed by direct enumeration methods, these results provide robust, comparative estimates of virion and virocell responses to wet-up and phosphorus availability. The opposing trends in virions and virocells provide direct evidence for a shift from environmental persistence to active infection following wet-up, reinforcing the interpretation that rewetting rapidly activates viral infection dynamics in soil ecosystems.

### Environmental DNA provided complementary evidence for rapid viral turnover following wet-up

eDNA can represent a substantial and temporally distinct nucleic acid pool that integrates over different time scales, relative to intact or transcriptionally active microbial and viral communities ^15,20^. eDNA represents a transient and independent pool that integrates viral turnover over short timescales, providing complementary insight into viral dynamics not captured by viromes or metatranscriptomes. We leveraged eDNA as an additional molecular fraction to contextualize viral dynamics following soil wet-up. Nucleic acid concentration measurements revealed pronounced and contrasting temporal responses across molecular pools (Fig. 5A). eDNA concentrations were highest in dry soils and declined sharply after wet-up, with only limited recovery over subsequent weeks, consistent with rapid turnover following rewetting. Elevated eDNA in dry soils is consistent with reduced microbial activity and decreased degradation or consumption of eDNA under dry conditions, as previously observed ^15,75^. The ∼2-fold decline immediately following wet-up likely reflects increased microbial activity ^22^, including uptake of eDNA as a nutrient source, as well as enhanced enzymatic degradation under moist conditions ^15,76^. Virome-derived DNA similarly decreased ∼2-fold after wet-up, likely due to a combination of virion degradation and increased host infection, but remained detectable throughout the experiment, whereas metatranscriptomic RNA concentrations were comparatively stable across time, reflecting persistent RNA or sustained microbial and viral transcriptional activity following rehydration (Fig 5A). Notably, eDNA concentrations increased by 1.4-fold from week one to two post–wet-up, coinciding with a continued decline in total soil DNA, presumably derived from microbial cells (Fig. 5A). This temporal correspondence is consistent with increased microbial turnover, potentially including viral-mediated lysis, contributing to eDNA pools during early post–wet-up succession. From weeks two to three, eDNA concentrations decreased again by ∼1.7-fold, alongside stabilization of viral and microbial community dynamics, suggesting the establishment of a more stable post-perturbation community. eDNA exhibited behavior distinct from both viromes and metatranscriptomes, consistent with a transient eDNA pool in dry soils that responds rapidly to wet-up and subsequent biological turnover (Fig. 5A).

**Figure 5:**
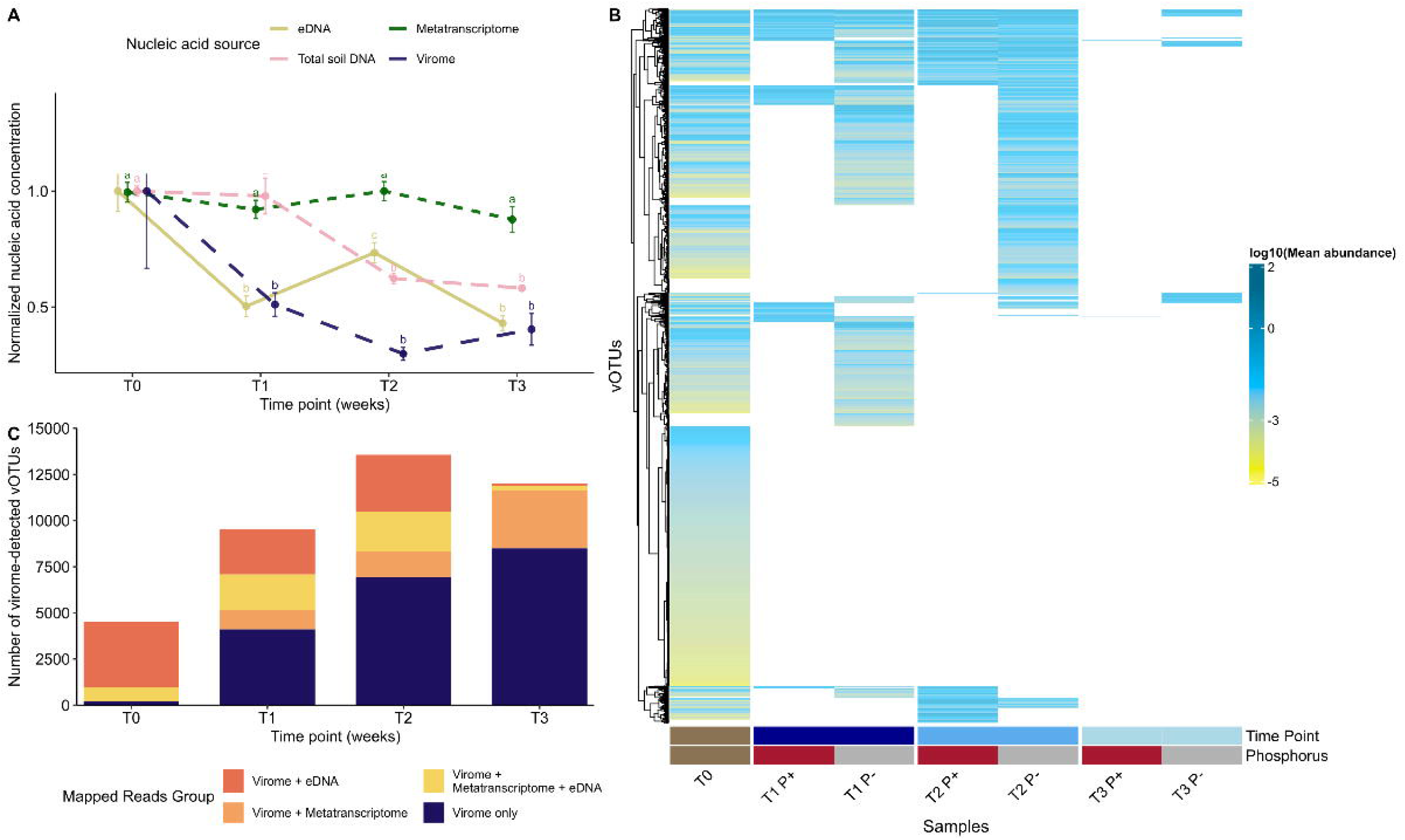
eDNA captures rapid and transient viral turnover following wet-up. (A) Mean normalized nucleic acid concentrations per gram of soil for environmental DNA (eDNA; yellow), metatranscriptomes (RNA; green), total soil DNA (pink), and viromes (blue) across dry soil (T0) and post–wet-up time points (T1–T3). Error bars indicate standard error. The left y-axis represents normalized nucleic acid concentrations where a value of one represents the time point with the highest mean nucleic acid concentration. Different letters denote statistically significant differences among time points within each nucleic acid source (P < 0.05), as determined by post hoc multiple-comparison tests (Tukey HSD or Dunn’s test, as appropriate); time points sharing a letter are not significantly different. (B) Heatmap showing log10-transformed mean normalized abundance of eDNA reads mapped to virome-derived vOTUs. Rows represent vOTUs and columns represent time points, with phosphorus treatment indicated by color bars below the heatmap. (C) Bar plot showing vOTUs detected in the virome at each time point and their simultaneous detection in eDNA and/or metatranscriptomes based on read mapping. Colors indicate the molecular fractions in which each vOTU was co-detected. T0 = dry soils; T1 = 1 week; T2 = 2 weeks; T3 = 3 weeks post–wet-up.

We mapped eDNA reads to virome-derived vOTUs and found that the viral eDNA signal was highest in dry soils and early post–wet-up time points. It became increasingly sparse and heterogeneous over time, with few vOTUs exhibiting persistent eDNA detection across multiple weeks (Fig. 5B). These results indicate that viral eDNA pools can turn over rapidly following perturbation and may not track either virion-associated DNA or transcriptionally active viruses on the same time scale. Notably, a greater number of unique vOTUs were detected in unamended soils across time points (Fig. 5B), potentially reflecting stronger nutrient limitation. Phosphorus-replete conditions may increase microbial growth ^77^ resulting in enhanced uptake and degradation of eDNA as a nutrient source ^76^, reducing the persistence of eDNA signals. Following wet-up, an increasing fraction of virome-detected vOTUs were co-detected in metatranscriptomes, while co-detection of eDNA in other fractions was not persistent over three weeks, with very few eDNA-detected vOTUs present by the third week (Fig. 5C). Importantly, these co-detection patterns reflect changes in detectability across molecular fractions and may or may not indicate transitions between viral states. Our results indicate that eDNA provides independent support for rapid viral turnover following wet-up and reinforces the interpretation that viromes, metatranscriptomes, and eDNA capture complementary but temporally distinct aspects of soil viral dynamics, further highlighting the importance of integrating multiple molecular fractions to resolve ecosystem-scale viral processes.

### Taxonomy-specific virus–host dynamics revealed responses of putative *Actinomycetota* phages to phosphorus amendment

In the 16S rRNA gene amplicon data, the two dominant bacterial phyla were *Actinomycetota* and *Pseudomonadota*, with relative abundances of 30% ± 4% and 25% ± 3%, respectively (Fig. 6A). Among all detected taxa, only the relative abundance of *Acidobacteriota* and *Myxococcota* responded significantly to phosphorus treatment (Dunn’s Test, *p* < 0.05). No statistically significant temporal changes in relative abundances were observed for bacteria at the phylum level (Fig. 6A). Analyses were conducted at the phylum level to match in silico virus–host predictions, as this level provides the highest confidence in virus–host assignments. The observed shifts in prokaryotic community composition and taxonomy-specific responses to phosphorus reflect life-history strategies aligned with prior studies. For instance, extractable phosphorus in soil along a phosphorus gradient explained a significant portion of variation in prokaryotic community composition ^28^, and bacterial community composition in grasslands was significantly impacted by phosphorus amendment ^78^. Lineage-specific responses to soil phosphorus concentrations have previously been observed ^28,78^, reflecting differences in microbial life history strategies (copiotroph or oligotroph). Our data support this hypothesis, as we also detected a significant decrease in relative abundance of *Acidobacteriota,* a phylum widely characterized to be oligotrophic ^79^, from 13.9% ± 1.22% to 11.0% ± 0.97% following phosphorus addition. There was also a significant positive correlation between the transcriptionally active viral and prokaryotic community structures (Spearman’s *r* = 0.24, *p* = 0.016; Supplementary Fig. 4A), suggesting that the transcriptionally active viruses were closely coupled to the prokaryotic community.

**Figure 6:**
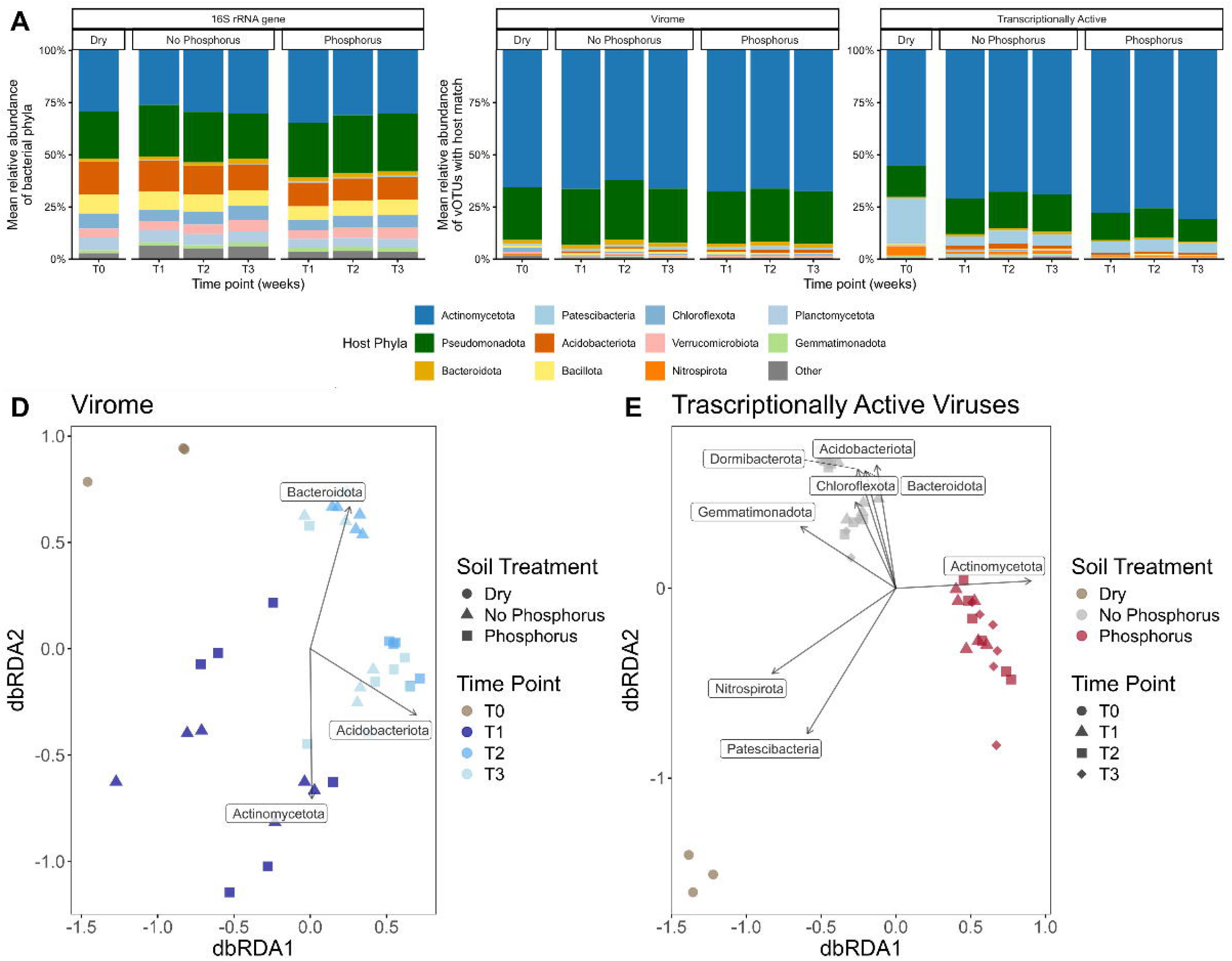
Virus–host interactions shaped by phosphorus amendment in transcriptionally active but not virion communities. Stacked bars show the relative abundances of phyla in the (A) 16S rRNA gene amplicon sequencing data or the relative abundance of vOTUs with host matches according to predicted host phylum in the (B) viromes or (C) transcriptionally active viral communities. Bars are colored by the 10 host phyla with the highest mean relative abundances in either viral group (virome or transcriptionally active), and all other phyla are grouped as “Other”. Distance-based redundancy analysis (dbRDA) plots show differences in viral community composition for comparing Bray-Curtis dissimilarity matrices of either the (D) viromes and (E) transcriptionally active viral communities to the sum relative abundance of viruses predicted to infect similar host (phylum). (D) Points are colored by time and shapes are soil type; (E) points are colored by soil type and shapes are time points. Vectors indicate the trajectories across viral community compositional space for the summed relative abundances of vOTUs by predicted host phylum for (D) viromes and (E) transcriptionally active viral communities. For viromes (D), only phyla that showed significant of the top 5 phyla based on overall relative abundance are shown. T0 = dry soil, T1 = 1 week, T2 = 2 weeks, T3 = 3 weeks post–wet-up.

To explore virus–host dynamics, we examined relative abundance patterns of viruses with a predicted host and host lineage-specific responses by using distance-based redundancy analysis (dbRDA). Among vOTUs with a predicted host, *Actinomycetota* and *Pseudomonadota* were the most abundant predicted host phyla across all time points and treatments, representing 66.0 ± 3.6% and 26.1 ± 2.6% of virion-associated vOTUs, and 72.3 ± 7.8% and 15.1 ± 4.1% of transcriptionally active viruses, respectively (Fig. 6B, C). The high relative abundance of these prokaryotic phyla and the viruses predicted to infect them aligns with previous studies in these HREC grassland soils, which also found high relative abundances of *Actinomycetota* and *Pseudomonadota* (formerly *Proteobacteria*) and/or their viruses ^15,23–25,80^. Although *Actinomycetota*-infecting phages were prevalent in both phosphorus-amended and unamended soils of transcriptionally active viruses (metatranscriptomics reads mapped to DNA vOTUs), their relative abundance significantly increased in the phosphorus-amended soils (78.1% ± 5.3%) compared to unamended soils (69.3% ± 3.2%; Dunn’s test, *p* < 0.001), while *Pseudomonadota-*infecting phages had higher relative abundance in the unamended soils (17.4% ± 2.5%; *p* < 0.05; Fig. 6B). This pattern was supported by dbRDA, where *Actinomycetota*-infecting phage loadings aligned with transcriptionally active viral communities in phosphorus-amended soils, indicating an association with phosphorus availability (Fig. 6E). Interestingly, neither *Actinomycetota* nor *Pseudomonadota* significantly increased in relative abundance in the phosphorus-amended soils, despite the increased relative abundances of their associated transcriptionally active viruses (Fig. 6B, C).

While other studies have not looked at viral activity in response to both soil rewetting and phosphorus amendment together, a recent study reported that a large proportion of transcriptionally active vOTUs were predicted to infect *Actinomycetota* and *Pseudomonadota* one week after wet-up ^63^. Additionally, phages predicted to infect *Actinomycetota* were previously shown to be enriched in dry soils ^15,16^, consistent with the high relative abundance of these phages in dry soils of the virome (65.6% ± 0.64%; Fig. 6B) and as transcriptionally active viruses (55.2% ± 1.3%; Fig. 6C) here. While the relative abundance of these predicted *Actinomycetota* phages remained relatively constant in the virome post-wet-up, their relative abundance significantly increased in the transcriptionally active viruses (74.4% ± 6.0%; Fig. 6D) following wet-up and remained consistently high throughout the three weeks, with 97% of *Actinomycetota* phage vOTUs initially detected in dry soils present at all post–wet-up time points (Supplementary Fig. 5A). This suggests that wet-up activates a persistent reservoir of virions, while phosphorus availability modulates subsequent infection dynamics by increasing viral transcription in *Actinomycetota* independently of host abundance.

## Conclusion

In a three-week grassland microcosm study, we used viromics, metatranscriptomics, and eDNA to track viral community dynamics following wet-up and phosphorus amendment. Viromes and transcriptionally active viruses represent distinct components of soil viral communities that respond differently to environmental perturbations. Rewetting of Mediterranean grassland soils following the dry season triggered a rapid increase in viral activity, reflected across molecular fractions with an increase in viral transcriptional activity and corresponding shifts in virion-associated populations. While wet-up may be the environmental change that initiates viral community activity, nutrient availability also plays a major role in infection and host dynamics. Phosphorus amendment increased the number of virocells overall, indicating enhanced viral infection, but also revealed decoupling between the relative abundance of transcriptionally active viruses and their predicted host phyla, particularly within *Actinomycetota*. This suggests that only a subpopulation of the highly abundant *Actinomycetota* is actively infected by viruses while other subpopulations are not, keeping the overall relative abundance stable.

By incorporating eDNA as a complementary molecular fraction, we provide independent evidence that eDNA pools respond rapidly to wet-up and integrate over shorter and distinct temporal scales relative to virion-associated DNA. Patterns observed in eDNA were consistent with rapid viral turnover following rewetting and reinforced the interpretation that different molecular fractions capture different and/or temporally offset aspects of soil viral dynamics. These findings support a model in which soil viral communities are structured by distinct but interacting molecular pools operating across different ecological timescales. Virions persist as a reservoir in dry soils, while rewetting triggers a transition to active infection, reshaping host populations and microbial turnover. Nutrient availability, particularly phosphorus, further regulates these dynamics by modulating infection intensity rather than virion abundance. This framework highlights viruses as active and responsive components of soil ecosystems that influence microbial community structure and biogeochemical processes following environmental perturbation. Incorporating these dynamics into ecosystem models will be critical for predicting microbial and biogeochemical responses to changing precipitation and nutrient regimes.

## Data availability

The sequences for the viromes are available on the JGI Gold database (Study ID # Gs0156767) and metatranscriptomics, eDNA, and 16S rRNA gene sequences are available at NCBI short read archive (SRA) NCBI under bioproject PRJNA1161162 (biosamples and SRA numbers available in Supplementary Table 16). All dereplicated vOTUs are available https://zenodo.org/records/19464936. R-code used to analyze data is provided on GitHub (https://github.com/gogogogogul/viral_activity_phosphorus).

## Author contributions

G.G. performed data curation, formal data analysis, visualization, writing and revising. G.M.A. assisted with experimental procedures and writing and editing. J.B.E and I.L. supervised data analyses, edited and revised the manuscript. S.J.B. and J.P.R. assisted with experimental procedures, supervised data analyses, edited and revised the manuscript. G.T. conceptualized the study, acquired funding, ran the experimental procedures, supervised, and revised the manuscript. All authors provided feedback on the final version.

## Acknowledgements

We acknowledge that HREC sits on the traditional, unceded land of the Pomo Indians. Research conducted at Lawrence Livermore National Laboratory took place on the territory of xučyun (Huichin), the ancestral and unceded land of the Chochenyo-speaking Ohlone people. Thanks to everyone in the Emerson lab for guidance and support. We thank Anneliek ter Horst and Hugo F. Monteiro for comments on a draft of the manuscript.

