## Supplemental Figures for "Divergent successional patterns and infection dynamics in virion and transcriptionally active soil viral communities following phosphorus amendment and wet-up"

Supplementary Figures and Captions

**Supplementary Figure 1: Shared vOTUs across treatments in the virome and transcriptionally active viral communities.** Venn diagrams show the number of vOTUs shared across treatments for (A) virome and (B) transcriptionally active viruses. Colors represent different treatments and percentages represent the proportion of shared vOTUs relative to the total number detected within each sample type.

**Supplementary Figure 2: Predicted virocell abundance across virus-per-cell estimates under phosphorus treatments.** Ribbon plots show predicted virocell abundances using varying viruses-per-cell ratios (indicated to the right of each line) for soils without phosphorus (left panels) and with phosphorus (right panels) across time points. Colored lines represent treatment means across time; ribbons indicate 95% confidence intervals. T0 = Dry Soil, T1 = 1 week, T2 = 2 weeks, T3 = 3 weeks post–wet-up. Temporal trends were similar across different number of viruses per infected cell (GAM followed by ANOVA, $p = 1$)

**Supplementary Figure 3: Shared vOTUs across time points in the virome and transcriptionally active viral communities.** Venn diagrams show the number of vOTUs shared across multiple time points for (A) virome and (B) transcriptionally active viruses. Colors represent different time points and percentages represent the proportion of shared vOTUs relative to the total number detected within each sample type.

**Supplementary Figure 4: Virome and microbial communities are closely coupled to the transcriptionally active viral community**. Pairwise Mantel tests (Spearman correlation) comparing community distance matrices derived from the virome, transcriptionally active viruses, and 16S rRNA gene amplicons. Each panel shows the relationship between distance matrices from two community types, indicated by the panel title. Each point represents the pairwise dissimilarity between two samples for one community type on the x-axis and another on the y-axis. Linear regression lines are shown only when the Mantel test is statistically significant ($p < 0.05$). Spearman correlation coefficient *p*-values are reported in the bottom right of each panel.

**Supplementary Figure 5: *Actinomycetota*-infecting phage of dry soils persist in post–wet-up time points.** Venn diagrams showing the number of *Actinomycetota* phage vOTUs shared across soil moisture (dry or wet) or time point for (A) virome and (B) transcriptionally active viruses

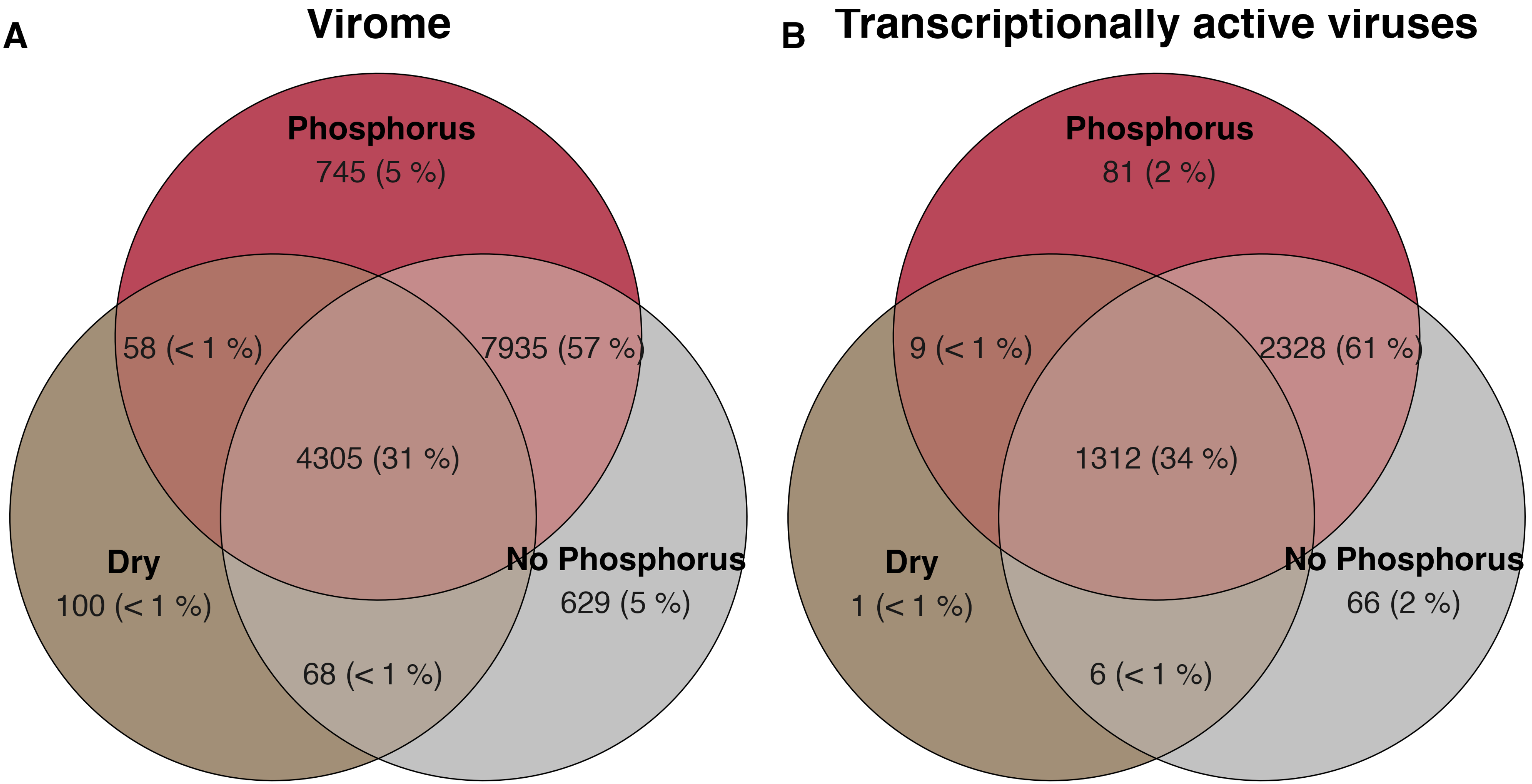
A
Virome
Phosphorus
745 (5 %)
58 (< 1 %)
7935 (57 %)
4305 (31 %)
Dry
100 (< 1 %)
No Phosphorus
629 (5 %)
68 (< 1 %)
B
Transcriptionally active viruses
Phosphorus
81 (2 %)
9 (< 1 %)
2328 (61 %)
1312 (34 %)
Dry
1 (< 1 %)
No Phosphorus
66 (2 %)
6 (< 1 %)

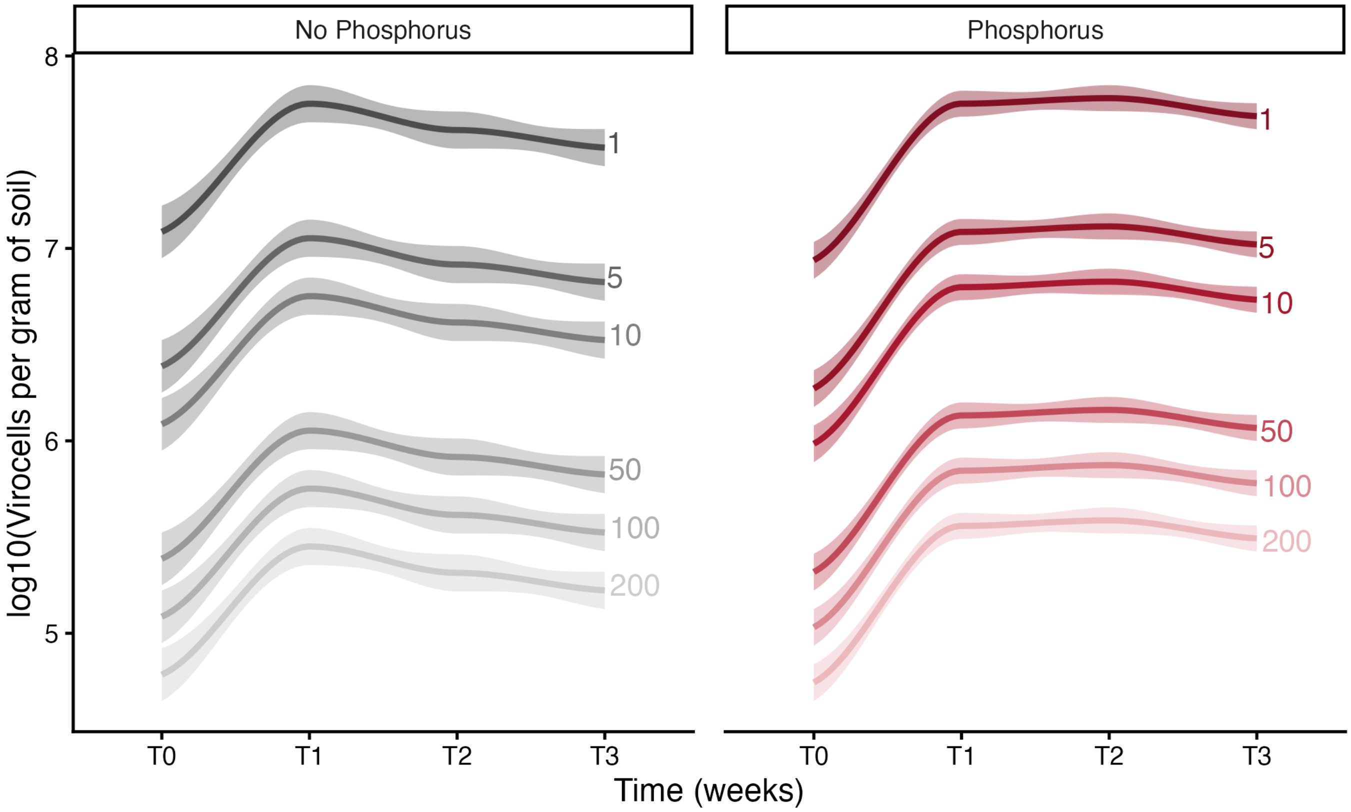
No Phosphorus
Phosphorus
log10(Virocells per gram of soil)
8
7
6
5
1
5
10
50
100
200
T0
T1
T2
T3
Time (weeks)

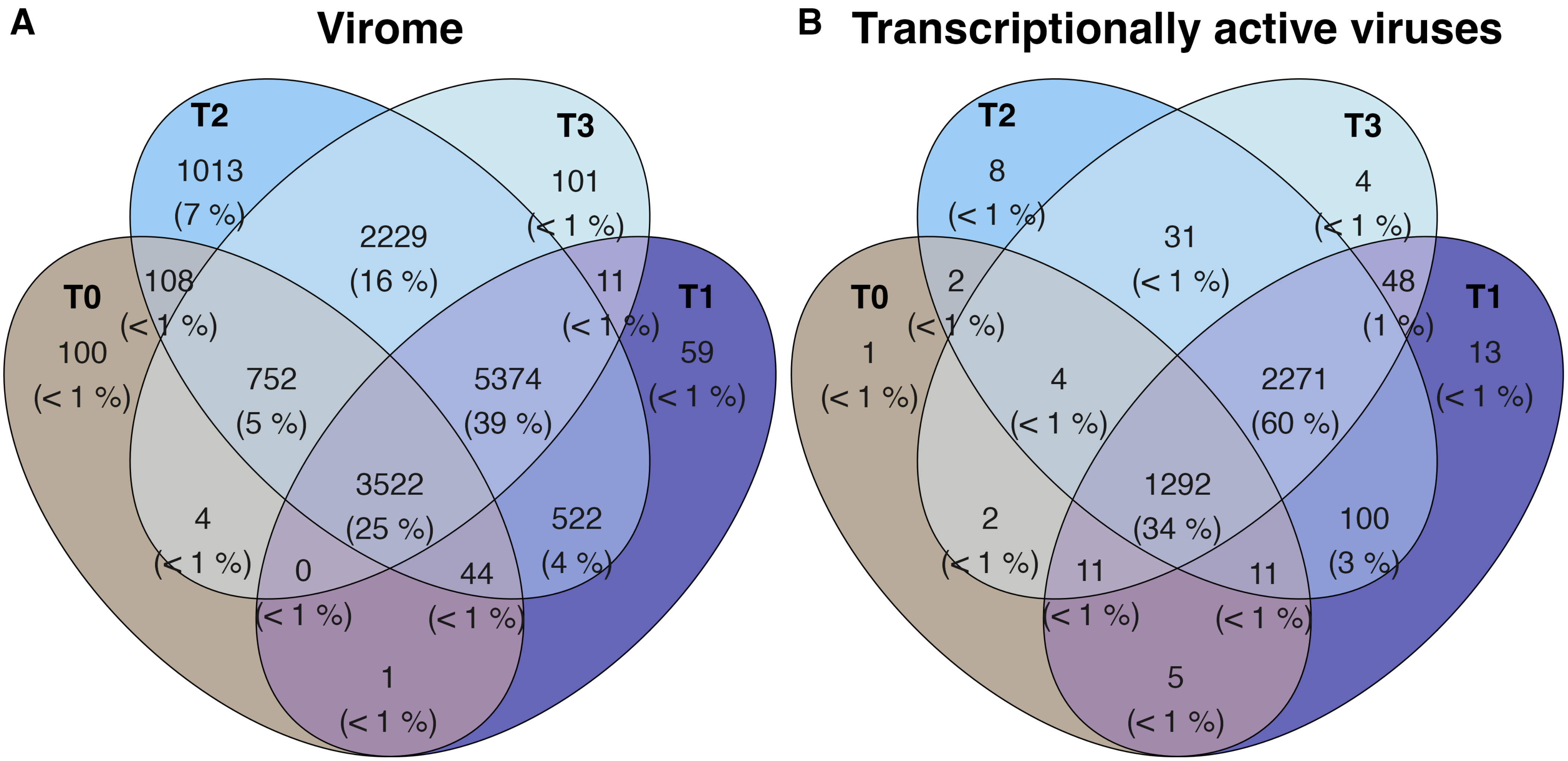

A
Virome
T2
T3
T0
T1
1013
(7 %)
2229
(16 %)
101
(< 1 %)
108
(< 1 %)
11
(< 1 %)
100
(< 1 %)
752
(5 %)
5374
(39 %)
59
(< 1 %)
3522
(25 %)
4
(< 1 %)
522
(4 %)
0
(< 1 %)
44
(< 1 %)
1
(< 1 %)
B
Transcriptionally active viruses
T2
T3
T0
T1
8
(< 1 %)
31
(< 1 %)
4
(< 1 %)
2
(< 1 %)
48
(1 %)
1
(< 1 %)
4
(< 1 %)
2271
(60 %)
13
(< 1 %)
1292
(34 %)
2
(< 1 %)
100
(3 %)
11
(< 1 %)
11
(< 1 %)
5
(< 1 %)

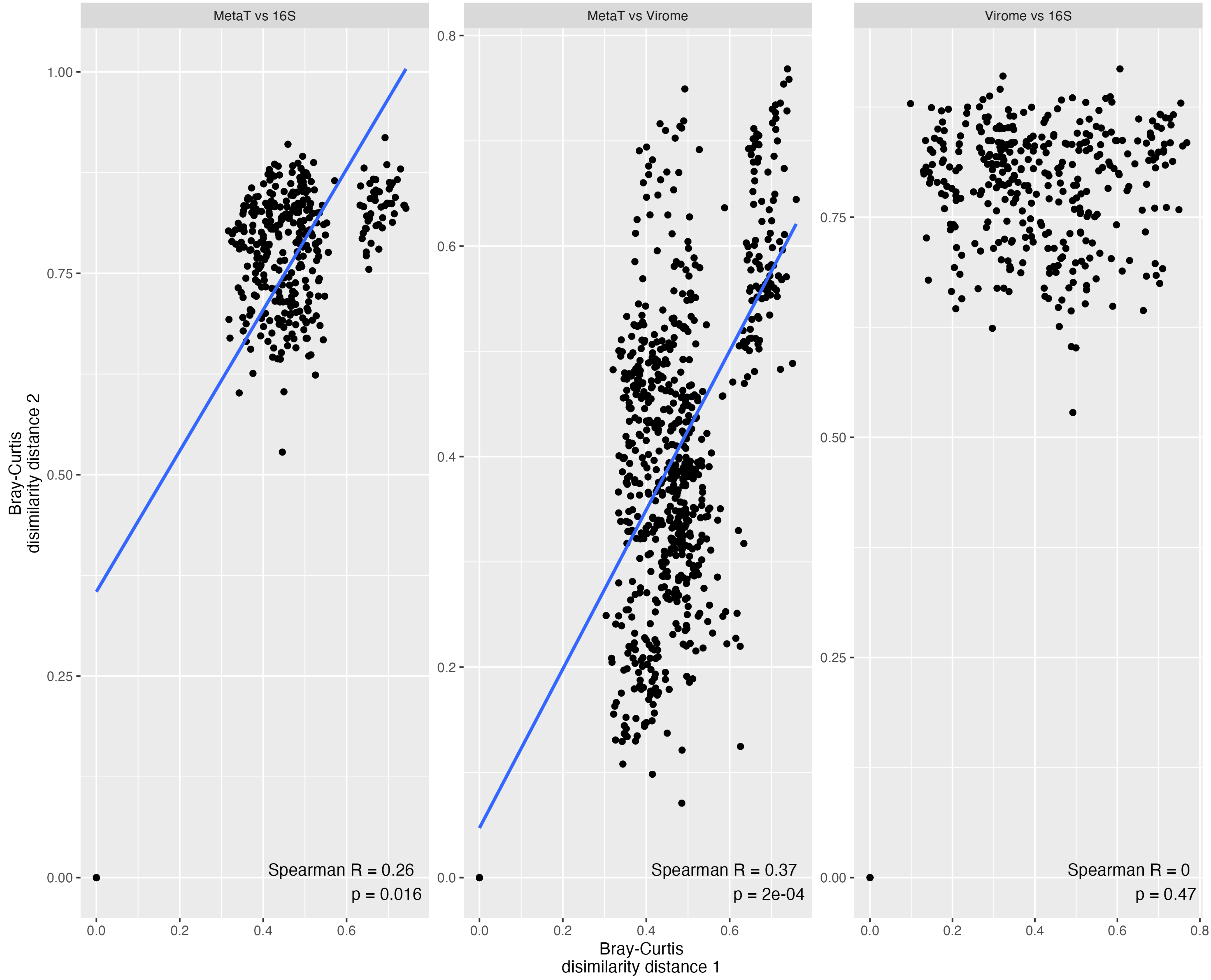

MetaT vs 16S
MetaT vs Virome
Virome vs 16S
Bray-Curtis
disimilarity distance 2
Bray-Curtis
disimilarity distance 1
Spearman R = 0.26
p = 0.016
Spearman R = 0.37
p = 2e-04
Spearman R = 0
p = 0.47

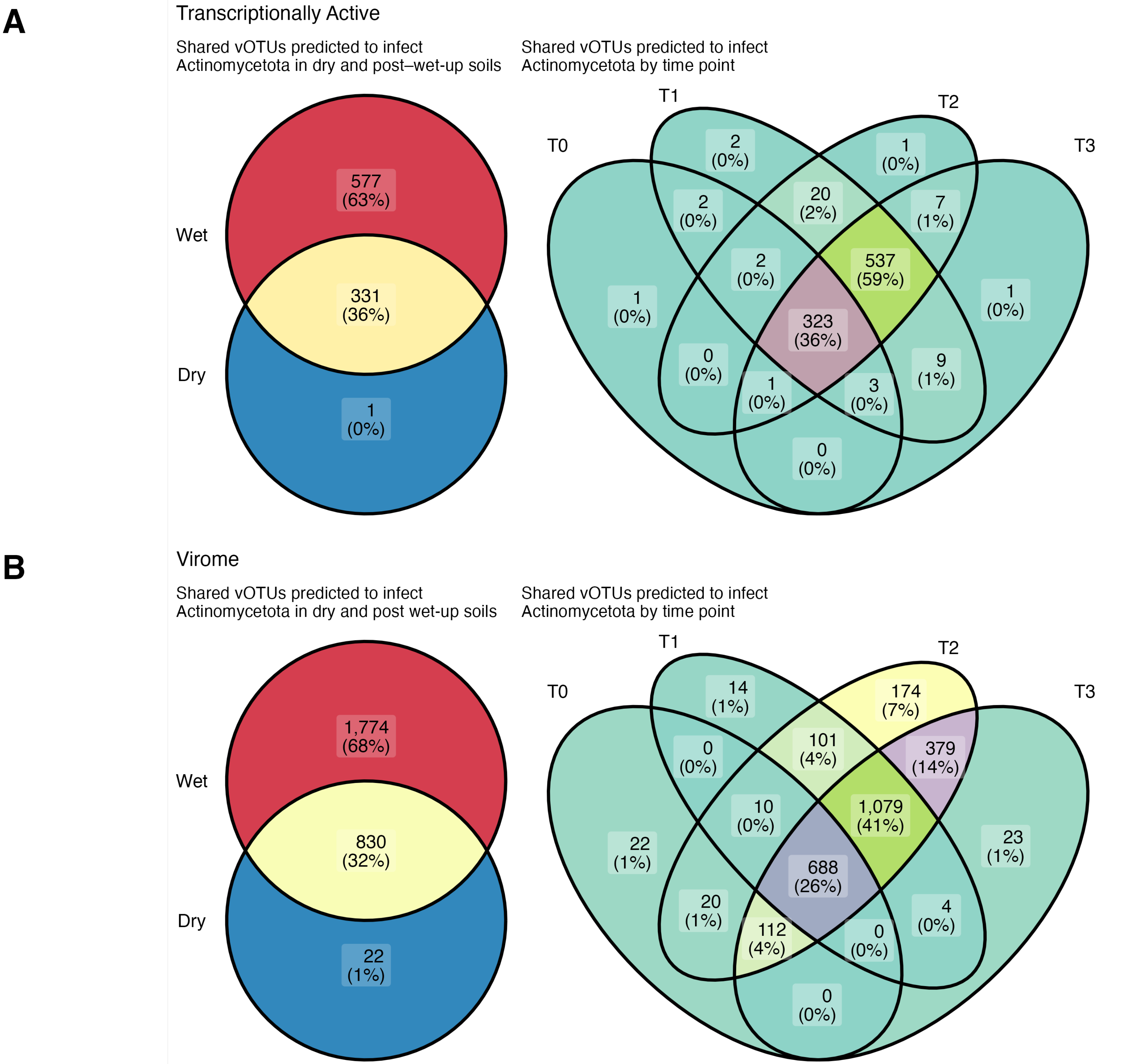
A
Transcriptionally Active
Shared vOTUs predicted to infect Actinomycetota in dry and post–wet-up soils
Wet
Dry
577 (63%)
331 (36%)
1 (0%)
Shared vOTUs predicted to infect Actinomycetota by time point
T0
T1
T2
T3
2 (0%)
1 (0%)
2 (0%)
20 (2%)
7 (1%)
2 (0%)
537 (59%)
1 (0%)
323 (36%)
1 (0%)
0 (0%)
9 (1%)
1 (0%)
3 (0%)
0 (0%)
B
Virome
Shared vOTUs predicted to infect Actinomycetota in dry and post wet-up soils
Wet
Dry
1,774 (68%)
830 (32%)
22 (1%)
Shared vOTUs predicted to infect Actinomycetota by time point
T0
T1
T2
T3
14 (1%)
174 (7%)
0 (0%)
101 (4%)
379 (14%)
10 (0%)
1,079 (41%)
22 (1%)
23 (1%)
688 (26%)
20 (1%)
4 (0%)
112 (4%)
0 (0%)
0 (0%)
