## Supplemental Methods for "Divergent successional patterns and infection dynamics in virion and transcriptionally active soil viral communities following phosphorus amendment and wet-up"

Supplementary Methods:

***Soil collection and experimental setup***

Soil was collected prior to the first seasonal rainfall in Autumn on October 3, 2020, from the Buck pasture plot at the Hopland Research and Extension Center (HREC; Mendocino County, CA; 39.001767° N, 123.069733° W) at an elevation of 1,066 feet and a depth of 0–10 cm. Soil was sieved using a 2.0 mm wire mesh, homogenized, and stored in large whirl pack bags and placed in a cooler at room temperature before being transported to Lawrence Livermore National Laboratory. Soil was collected for gravimetric water content (GWC), pH measurements, and placed in microcosms.

Microcosms were established by adding 206 g (± 0.5 g) of soil to 1.9 L acid-washed and autoclaved Mason jars. Microcosms received one of two treatments: (1) deionized water (dH_2_O), pH 5.5 (natural pH of soil at sampling site (Foley et al., 2023); or (2) dH_2_O, pH 5.5 + KH_2_PO. Water was evenly applied to the soil using a syringe to achieve 30% GWC. Treatments were replicated six times for each of the three time points (days 7, 14, and 21 post wet-up), totaling 36 microcosms (two treatments, six replicates, three time points). For phosphorus amendment treatment, KH_2_PO_4_ was added to water, for a final concentration of 50 µM (1.37 μg P/g soil), increasing phosphorus availability by approximately 6–9% above the baseline concentrations previously recorded at this location (Eviner et al., 2006; Nuccio et al., 2016). Mason jars were sealed with lids containing airtight septa and incubated at room temperature (~23°C).

Both the initial dry soil and soils collected from microcosms were sampled (39 total samples; 3 dry soils and 36 from microcosms), with the microcosms destructively sampled on days 7 (T1), 14 (T2), and 21 (T3) post wet-up. The wet soil was homogenized, and 5 g was collected for GWC measurements, 10 g for pH measurement, 0.5 g was collected into 2.0 mL Lysing Matrix E tubes (MP Biomedicals) for DNA/RNA extractions (Barnard et al., 2013) for metagenome (not used in this study) and metatranscriptomes, and 10 g for virome processing. GWC and pH measurements (see supplementary methods), and DNA/RNA extractions were immediately conducted, and virome soil was frozen at –80 °C until processing the next morning.

***Soil gravimetric water content and pH measurements***

GWC measurements were previously reported (Sieradzki et al., 2025). Briefly, 5 g of soil in duplicate from dry soils and the microcosms from the last two weeks (two and three) were dried to a constant weight at 105 °C for 48 hours (Supplementary Table 2). The GWC of initial soils were back calculated for an initial GWC of 8.3%. We used our target GWC of 30% for the first week.

Soil pH was determined by adding 25 mL of deionized water (dH2O) to 10 g of soil, shaking for 1 hour at 400 rpm, and then allowing it to rest for 1 hour before measuring with a sensION+ pH meter (Hach) (FAO, 2021).

***Soil virome processing and DNA extraction***

The 10 g of soil that were collected and stored at –80°C for viromes were thawed at 4°C for 1 hour prior to viral like particle (VLP) suspension following previously established protocols (Trubl et al., 2019, 2016) with some minor amendments. Soil was washed three times with a total of 50 mL AKC’ buffer (1% w/v Potassium Citrate (K_3_C_6_H_5_O_7_), 10% v/v 10x Phosphate Buffered Saline (PBS), pH 7.4, 150 mM MgSO_4_, 50 µM Na_2_HPO_4_, 20 µM KH_2_PO_4_). The soil was shaken horizontally for 15 minutes at 400 RPM followed by three cycles of alternating vortexing and hand shaking for 30 seconds each, then centrifuged at 4 °C, 4,198 x *g* for 10 minutes. Approximately 50 mL of wash fluid was collected and passed through a Steriflip-GP 0.22 µm filter (Millipore Sigma; cat. # SCGP00525) to remove most cellular debris, the resulting filtrate was then concentrated to 250 µL using a 100 kDa Amicon Ultra-15 cellulose membrane filter (Millipore Sigma; cat. # UFC910096) that was pretreated with 3 mL of 1x PBS with 0.05% Tween 20 pH 7.4 (Teknova; cat. # P0201). Additionally, the Amicon filter was washed three times with 750 µl of AKC’ buffer per wash. The resulting 2.5 mL of VLP concentrate was then DNase treated with 7.1 µL of RQ1 RNase-Free DNase (Promega; cat. # M6101) and 270 µL DNase Reaction Buffer (0.1 M Tris-HCL pH 7.5, 25 mM MgCl_2_, 5 mM CaCl_2_) for 30 minutes at 37 °C to remove DNA not encapsidated by a VLP. The reaction was stopped by adding 135 µL of DNase Stop Buffer (Promega; cat. # M6101) incubated at 65 °C for 10 min, then stored overnight at 4 °C.

VLPs were then precipitated by adding 0.5 mM FeCl_3_ to each sample, and DNA was extracted following previous methods (Griffiths et al., 2000; John et al., 2011). Samples were centrifuged at 4 °C, 16,000 x *g* for 30 minutes. Pellets were resuspended in 20 µL of 0.2 M EDTA and 0.4 M ascorbic acid (~pH 6), and then 230 µL 1x TE buffer (pH 8) and 250 µL phenol/chloroform/isoamyl alcohol 25:24:1 (pH 8) was mixed in. Samples were incubated on ice for 20 minutes, vortexed intermittently to homogenize, and centrifuged at 4 °C at 16,000 x *g* for 5 minutes. The aqueous phase was then transferred to Phase-Lock Gel tubes (QuantaBio, Cat. #2302830) followed by adding 250 µL of chloroform, hand mixing and centrifuged at 16,000 x *g* for 5 minutes at 4 °C. The aqueous phase was transferred to a 1.5 ml microcentrifuge tube, where 25 µL of 3 M sodium acetate (pH 5.5), 1.5 µL Glycoblue (ThermoFisher, Cat. #AM9516), and 250 µL isopropanol were added to precipitate the DNA. Samples were incubated at least 1 hour at -20 °C followed by centrifuging samples at 16,000 x *g* for 20 minutes at 4 °C. Supernatant was discarded, and the resulting pellets were washed with 70% ethanol. Samples were centrifuged at 16,000 x *g* for 5 minutes at 4 °C, and the supernatant was discarded. DNA was then resuspended in 50 µL of 10 mM Tris-HCl pH 8. The resulting DNA was quantified using Qubit 2.0 fluorometer (Invitrogen; Supplementary Table 2), and purity was assessed with a NanoDrop 200 spectrophotometer (ThermoFisher), and the DNA was stored at –80 °C.

DNA was shipped to the Joint Genome Institute (JGI; Berkeley, CA) for library preparation and Illumina sequencing. Initially, DNA libraries were prepared for 16 of the 39 samples (Supplementary Table 1) using the Accel-NGS 1S kit (Swift Biosciences) and sequenced using the Illumina NovaSeq platform with 2x151bp indexed sequencing. Approximately 300M paired-end reads were generated per sample. The Accel-NGS 1S kit was chosen due to its previously verified capability to quantitatively amplify both single-stranded and double-stranded DNA (Roux et al., 2017; Trubl et al., 2025, 2019). Due to availability issues of the Accel-NGS 1S kit, the previously tested SRSLY NGS library Prep Kit (Claret Biosciences, Santa Cruz, CA, USA) was used to redo 38 of the 39 samples. One replicate from week three was not used due to insufficient DNA (T3H218O_A). These libraries were also sequenced using the Illumina NovaSeq platform with 2x151bp indexed sequencing, again producing approximately 300M paired-end reads per sample. Together, the two library preparation methods produced a total of 54 viromes (see Supplementary Table 1).

***Bulk soil RNA extraction and read processing***

RNA was extracted following previously published methods (Barnard et al., 2013), briefly, soil was combined with 200 µl of TE (pH 8.0), 500 µl of extraction buffer (5% CTAB, 0.7M NaCl, 240mM KPO_4_, pH 8) and 500 µl of ice-cold 25:24:1 phenol:chloroform:isoamyl alcohol in tube. Tubes were shaken in a FastPrep (MP Biomedicals) for 30 seconds at speed 5.5 m/s, spun at 16,000 x *g* at 4 °C, and then the aqueous layer was transferred to a 2.0 ml Phase-Lock gel tube (QuantaBio Cat. #2302830). This process was repeated on the organic layer and 500 µl 24:1 chloroform:isoamyl alcohol was added to the aqueous layer and spun at 16,000 x *g* at 4 °C. The aqueous layer was added to a 2.0 ml microcentrifuge tube with 1 ml 40% PEG6000, 1.6M NaCl, and 1 µl GlycoBlue (ThermoFisher Cat. AM9516) and incubated overnight at room temperature. Samples were precipitated and resuspended in 50 µl of TE (pH 8.0). The RNA fraction was cleaned up using the Qiagen All-Prep kit (Cat. #80284), following the manufacturer's protocol. RNA library preparation and sequencing methods were previously reported (Sieradzki et al., 2025). Briefly, RNA samples were treated with RQ1 RNase-Free DNase (Promega Cat. # 89836), then sent to Azenta for library preparation with the NEBNext Ultra II RNA Library Prep Kit (NEB, Ipswich, MA, USA), rRNA depletion with the Fast select rRNA depletion Kit (Bacteria). The samples were sequenced using an Illumina HiSeq 2500 instrument according to manufacturer’s instructions using a 2x150 Paired End (PE) configuration, resulting in ~350 M raw paired end reads (~105 GB sequencing).

Reads were checked for quality and trimmed by first trimming to remove any contamination, including barcodes, primers, and adapters using BBDuk (v. 38.63; Bushnell, 2014) (t=24, ftl=5, ktrim=r, k=23, mink=11, hdist=1, tbo, tpe, minlen=50), then reads were filtered for any remaining barcodes, primers, and PhiX , where the read and its pair were removed if a match was found (k=31, hdist=1, minlen=50). Finally, low-quality sequences were removed from reads (qtrim=r, trimq=10, minlen=50).

***Environmental DNA extraction, sequencing, and analysis***

Environmental DNA (eDNA) was extracted from 5 g of soil thawed at 4 °C for 1 hour. Soil was washed twice with a total of 25 mL of 10 mM Tris-HCl (pH 8.0) by shaking horizontally for 15 minutes at 400 RPM at room temperature, followed by three cycles of alternating vortexing and hand mixing for 30 s each. Samples were then centrifuged at 4 °C at 4,198 × *g* for 20 min. Approximately 23 mL of wash fluid was collected and passed through a 0.22 µm Steriflip-GP filter (Millipore Sigma; cat. # SCGP00525) to remove cellular debris, followed by filtration through a 0.02 µm filter (Whatman Anotop, Sigma Aldrich; cat. # WHA68091002) to remove viral particles.

The resulting filtrate was concentrated by isopropanol precipitation as described above. Briefly, approximately 20–25 mL of filtrate was amended with 23 µL GlycoBlue (Thermo Fisher Scientific), 2.3 mL of 3 M sodium acetate, and 23 mL of isopropanol, mixed thoroughly, and incubated at −20 °C for at least 1 hour. Samples were centrifuged at 4,198 × *g* for 20 minutes at 4 °C, and the resulting DNA pellet was washed with freshly prepared cold 70% ethanol. After a final centrifugation (5 min at 4,198 × *g*, 4 °C) and removal of residual ethanol, pellets were air-dried for approximately 30 min and resuspended in 100 µL of 10 mM Tris-HCl (pH 8.0). DNA concentrations were quantified using a Qubit 2.0 fluorometer (Invitrogen; Supplementary Table 2), and samples were stored at −80 °C until sequencing.

eDNA samples were sent to GENEWIZ, Inc. (South Plainfield, NJ, USA) for library preparation and sequencing. DNA libraries were prepared using the NEBNext Ultra DNA Library Preparation Kit (New England Biolabs, Ipswich, MA, USA) following the manufacturer’s recommendations. Briefly, eDNA was fragmented by acoustic shearing using a Covaris S220 instrument. Fragmented DNA was end-repaired and adenylated, followed by ligation of sequencing adapters. Adapter-ligated DNA was indexed and enriched by limited-cycle PCR (8–17 cycles). Final libraries were assessed for fragment size distribution using a TapeStation system (Agilent Technologies) and quantified using a Qubit 2.0 fluorometer. Libraries were further quantified by quantitative PCR, pooled, and sequenced on an Illumina HiSeq platform using a 2×150 bp paired-end configuration.

Raw eDNA reads were processed in three steps using BBDuk (v. 38.63; Bushnell, 2014). First, sequencing adapters were trimmed using TruSeq adapter references (ktrim=r, k=23, mink=11, hdist=1, tbo, tpe, minlen=50). Second, PhiX contamination was removed (k=31, hdist=1, minlen=50). Third, reads were quality trimmed (qtrim=r, trimq=10, minlen=50). Quality-controlled reads were mapped to viral operational taxonomic units (vOTUs) identified from viromes using Bowtie2 (v. 2.5.2; Langmead and Salzberg, 2012) with the sensitive alignment setting. Resulting SAM files were converted to BAM format, sorted, and indexed using SAMtools (v. 1.19; Danecek et al., 2021; Li et al., 2009). vOTU abundance tables were generated using CoverM (v. 0.6.1; Aroney et al., 2025) with the contig setting and both mean and count methods, requiring a minimum of 90% read identity. Using the resulting count tables, vOTUs were considered detected in a sample only if the summed number of reads mapped across triplicate samples exceeded three reads. The resulting mean coverage tables were filtered to include only vOTUs detected in each sample and normalized by total gigabases (Gbp) sequenced per eDNA library. Final abundance tables represent vOTU relative abundances normalized by contig length and sequencing depth (Supplementary Table 3).

eDNA and other nucleic acid concentrations were normalized to the highest mean value across time points (initial soils, T1, T2, or T3), such that a value of 1 represents the time point with the highest mean nucleic acid concentrations for each source.

***Virome read processing and vOTU identification***

For samples prepared using the Swift Accel-NGS 1S library preparation kit, raw reads were processed in three steps using BBDuk (v. 38.63; Bushnell, 2014). First, adapters were trimmed using TruSeq adapters references (ktrim=r, k=23, mink=11, hdist=1, tbo, tpe, minlen=50). Second, PhiX was removed (k=31, hdist=1, minlen=50). Third, reads were quality trimmed (qtrim=r trimq=10 minlen=50) and based on manufactures recommendations the last 10 base pairs on the forwarded reads (ftr=140) and first 10 base pairs of the reverse reads were trimmed (ftl=10). The resulting quality controlled reads were assembled using SPAdes (v. 3.15.4; Bankevich et al., 2012; Nurk et al., 2013) single cell module (-sc) with a kmer size (-k) of 21, 33, 55, 77, and 99 (Trubl et al., 2019). For samples sequenced using the SRSLY library kit, the same quality control steps were used as mentioned above, except the last 10 base pairs on the forwarded reads and first 10 base pairs of the reverse reads were not trimmed.

Viral contigs were identified from the resulting contigs using VirSorter2 (v. 2.2.4; Guo et al., 2021) and geNomad (v. 1.7.1; Camargo et al., 2023). First, viral contigs were identified using VirSorter2 following a pre-established protocol that involved an initial pass of VirSorter2(– virsorter run –keep-original-seq include-groups dsDNAphage, NCLDV, RNA, ssDNA, lavidaviridae --min-score 0.5 --min-length 5000). Next, viral contigs were assessed with CheckV (v.1.0.1; Nayfach et al., 2021) and retained following previously established criteria (Guo et al. 2021). Briefly, a predicted viral contig was kept if it met any one of the following: (i) contained ≥ 1 viral gene; (ii) contained 0 viral genes and 0 host genes; (iii) contained 0 viral genes and had a viral score ≥ 0.95; (iv) contained 0 viral genes and more than 2 hallmark genes; or (v) contained 0 viral genes, 1 host gene, and was > 10 kbp in length. The virus and provirus fasta files were then combined and passed through VirSorter2 once more (--seqname-suffix-off, --viral-gene-enrich-off, --provirus-off, --prep-for-dramv, --include-groups dsDNAphage, NCLDV, RNA, ssDNA, lavidaviridae, --min-score 0.5, --min-length 5000). The contigs identified as viral for geNomad were confirmed using CheckV and retained based on similar criteria as above: (i) contained ≥ 1 viral gene; (ii) contained 0 viral genes and 0 host genes; (iii) contained 0 viral genes and > 2 hallmark genes. The viral contigs that were identified by both VirSorter2 and geNomad were separated into two groups based on length to capture ssDNA and dsDNA viruses. The first group for dsDNA viruses consisted of viral contigs that were ≥ 10 kbp and the second for ssDNA viruses was ≥ 1kbp and < 10 kbp. For viral contigs between 1kbp and 10 kbp (identified only by geNomad), only those that were identified as *Monodnaviria* by geNomad were retained. The resulting viral contigs were combined and clustered, using previously established parameters (>95% ANI, over 85% of the contig) to generate a dereplicated set of viral populations (vOTUs) (Roux et al., 2019).

***vOTU relative abundances in viromes and metatranscriptomes***

Quality trimmed reads from the SRSLY library preparation kit were mapped to vOTUs, using for Bowtie2 (v. 2.5.2; Langmead and Salzberg, 2012) with the sensitive setting. The resulting SAM files were converted to BAM files, sorted (samtools sort), and indexed (samtools index) using SAMtools (v. 1.19; Danecek et al., 2021; Li et al., 2009). vOTU abundance tables were generated with CoverM (v. 0.6.1; Aroney et al., 2025), with the contig setting and trimmed_mean, applying previously established thresholds of >90% read identity covering at least 75% of the contig (Roux et al., 2017; Trubl et al., 2018). Abundance tables were then normalized by total Gbp of each virome producing relative abundances normalized by contig length and sequencing depth (Supplementary Table 4).

Quality-processed metatranscriptomic reads were mapped using Bowtie2 sensitive setting to DNA vOTUs and genes prediction and annotation using Prodigal-gv (v.2.11.0-gv; Camargo et al., 2023; Hyatt et al., 2010). SAM files were converted to BAM files, sorted, and indexed as described above. To identify transcriptionally active vOTUs, parameters similar to those previously described were applied (Emerson et al., 2018; Muscatt et al., 2022). Specifically, coverage tables were generated for metatranscriptomic reads mapped to vOTUs, then CoverM (v. 0.6.1; Aroney et al., 2025) was run with the contig setting and mean method (-m mean), requiring a minimum of 90% read identity. To generate a count table of reads mapped to predicted genes identified by Prodigal-gv (v.2.11.0-gv; Camargo et al., 2023; Hyatt et al., 2010), CoverM was run with the count method (-m count), also requiring at least 90% read identity (Muscatt et al., 2022). For each sample, if the total number of reads mapped to predicted genes across triplicate metatranscriptomes was fewer than four, the abundance of that gene for that sample was set to zero (Muscatt et al., 2022). A vOTU, including vOTUs that were identified as ssDNA viruses, were considered transcriptionally active only if at least one gene was considered expressed per 10 kbp of the vOTU across the data set, and vOTUs not meeting this criterion were assigned an ‘active’ relative abundance of 0 (Emerson et al., 2018; Muscatt et al., 2022). The mean coverage table of the transcriptionally active vOTUs was normalized by library size (Gbp) per metatranscriptome, generating the abundance table used for downstream analyses (Supplementary Table 5).

***Auxiliary metabolic genes identification***

Auxiliary metabolic genes were identified using DRAM-v (v 1.4.6; Shaffer et al., 2020). To generate DRAM-v compatible inputs, all vOTUs were process with Virsorter2 using minimum parameters (-seqname-suffix-off --viral-gene-enrich-off --provirus-off --prep-for-dramv --include-groups dsDNAphage,NCLDV,RNA,ssDNA,lavidaviridae --min-length 0 --min-score 0 all). DRAM-v was run with the skip tRNA scan and a minimum contig size of 1000 (--skip_trnascan --min_contig_size 1000). Resulting output was filtered for an auxiliary score less than four and did not contain an A (when the gene has been given identifiers associated with viral host attachment and entry), T (transposon nearby), or V (gene is assigned a VOGDB identifier with the replication or structure category) flag. Supplementary Tables 6–7 provides all genes identified by DRAM-v and a list of filtered AMGs.

***Host prediction***

Virus hosts were predicted using iPHoP (v1.3.3; Roux et al., 2023) with the iPHoP_db_Aug23_rw database (iphop predict –min_score 75 –db_dir Aug_2023_pub_rw). Genus-level host predictions were only reported when the confidence score was ≥ 90%, otherwise predictions at the family or higher taxonomic level were used (Trubl et al., 2025). First, normalized count tables were filtered to retain only vOTUs with a detectable host prediction, then relative abundances of each vOTU were calculated within each sample, and samples and vOTUs were removed if they either had zero vOTUs (samples) or had relative abundances of zero in every sample (vOTUs). Average vOTU relative abundances within host phyla were calculated and the top 10 phyla for each data type (viromic and metatranscriptomics) were retained while the remainder of vOTUs with a predicted host were changed to the “Other” category, and proportional abundances were then calculated and plotted.

***Amplicon processing***

DNA was extracted from 0.5 g of soil using the Powersoil Pro kit (Qiagen, Hilden, Germany) following manufacturer’s protocol. Library preparation and sequencing methods were previously published for 16S rRNA gene amplicon sequencing (Sieradzki et al., 2025). Briefly the hypervariable region v4 of the 16S rRNA gene was amplified using primers 515F–Y (515F-GTGYCAGCMGCCGCGGTAA) (Parada et al., 2016) and 806R (806R-GGACTACNVGGGTWTCTAAT) (Apprill et al., 2015). PCR reactions and subsequent PCR clean-up steps were performed as described by Sieradzki et al. (2025). The resulting amplicons libraries were then sequenced on the Illumina MiSeq platform using a 2x250 cycle kit with approximately 25% phiX spike-in.

Raw reads were examined for sequence quality, using previously published methods (Foley et al., 2023). Briefly, using BBDuk (v. 38.63; Bushnell, 2014) in sequential steps, adapters were trimmed (ktrim=r k=31 mink=11 hdist=1 tpe tbo) and PhiX was removed (k=31 hdist=1 ftl=10). Resulting reads were further scrutinized using DADA2 (v.1.30.0; Callahan et al., 2016), using the filterAndTrim function (truncLen=c(240,200), maxN=0, maxEE=c(2,2), truncQ=2, rm.phix=TRUE, compress=TRUE, multithread=FALSE), error rates were determined using learnErrors functions and corrected. The dada function of DADA2 was used on the full dataset using the inferred error, pair-end reads were merged with mergePairs and a sequence table was generated using makeSequenceTable. Chimeras were then removed from the generated sequence table. Taxonomy was assigned to each ASV using DADA2’s assignTaxonomy functions with the GTDB_bac120_arc53_ssu_r214_genus reference database. ASVs lacking a kingdom-level assignment were removed from the final abundance table.

Amplicon abundance tables for prokaryotic communities were generated by rarefying each sample to the 10th percentile of read depth specific to each amplicon type. Three samples from the prokaryotic (T1H218O_A, T1H2O_A, T1H2OP_C) communities were removed due to insufficient sequencing depth (Supplementary Table 9).

The predictions of virion and virocell abundances were performed similarly to prior work (Nicolas et al., 2023). To predict virion abundance, the fraction of viromic reads mapped to vOTUs was multiplied by the total amount of DNA extracted to estimate total viral DNA (equation 1), for each replicate. The mean of the weight of all vOTUs within a treatment type (phosphorus or without phosphorus) was obtained using a custom script considering the sum weight of each individual nucleotide, multiplied by 2 for viruses not identified as a ssDNA virus. Virion counts were predicted using equation 2. The number of virions for each sample was normalized based on the amount of dry soil used for virome (DNA) extractions.

${DNA}_{viral}(ng)= DNA (ng) \times\frac{Number of reads mapped to vOTUs per sample}{Total number of reads in sample}$ ( 1 )

$Particles (virions)=\frac{{DNA}_{viral} (ng)\times6.0221\times{10}^{23}(\frac{particles}{mol})}{{DNA}_{weight}\times1\times{10}^{9}(\frac{ng}{g})}$ ( 2 )

To predict virocell abundance, for each sample, metatranscriptomics reads that mapped to a DNA vOTU were considered viral reads, which was then divided by the total metatranscriptomics reads of the sample, to obtain a proportion of viral metatranscriptome reads per sample. The viral proportion was multiplied by the total amount of RNA extracted to estimate total viral RNA (equation 3). The mean weight of all vOTUs within a treatment type (phosphorus or without phosphorus) was obtained as previously described. Virus abundances were predicted using equation 4, which uses the total weight of viral RNA divided by the average vOTU weight to obtain moles of RNA viruses, this is then multiplied by Avogadro’s number to obtain number of viruses. At the viral community level, the number of viruses produced per infected cell (burst size) can vary, so a range of viruses per cell was used, from 1 virus per cell to 200 viruses per cell, but similar trends were observed across the range (ANOVA, *p* = 1; Supplementary Fig. 2; equation 5). To normalize based on the amount of soil used for metatranscriptome (RNA) extractions, the number of virocells for each sample was divided by the corrected dry weight for each extraction.

***Statistical analyses***

All statistical analyses were conducted in R (v.4.4.2; R Core Team, 2024) and rstatix (v.0.7.2; Kassambara, 2023). Viral community beta-diversity was assessed by computing Bray-Curtis dissimilarity matrices on Hellinger transformed vOTU abundance tables, using vegan (v.2.6-10; Oksanen et al., 2025) and ape (v.5.8-1; Paradis and Schliep, 2019). PERMANOVA tests were performed on the Bray-Curtis dissimilarity matrices, using the adonis2 function from vegan (v.2.6-10; Oksanen et al., 2025). Pairwise PERMANOVA comparisons were conducted on Bray-Curtis dissimilarity matrixes, using the pairwise.adonis function from the pairwiseAdonis package (v.0.4.1; Arbizu, 2017).

Host-virus interaction analyses were conducted using distance-based redundancy analysis (dbRDA) on Bray-Curtis dissimilarity matrices of Hellinger transformed vOTU abundance tables using the dbrda function in vegan. Significance of explanatory variables was tested using permutation-based ANOVA (anova.cca).

Differences in virion and virocell abundances across weeks and phosphorus treatments were assesed using ANOVA on linear models for each variable (week or phosphorus amendment), followed by Tukey’s HSD post hoc tests. To evaluate whether virocell estimates exhibited consistent temporal trends across assumptions of viruses per infected cell, generalized additive models (GAMs) were fitted using log10-transformed virocell abundance per gram of soil as a function of the interaction between viruses per cell and time point (treated as a categorical variable).

To determine changes in nucleic acid concentration over time, linear models were fitted separately for each nucleic acid source using normalized nucleic acid concentration per gram of soil modeled as a function of time point. Normality of residuals was assessed using Shapiro-Wilk test. For datasets meeting normality assumptions (virome, metatranscriptome, and eDNA), differences among time points were assessed using ANOVA, followed by Tukey’s HSD post hoc tests. For total soil DNA, which did not meet normality assumptions, differences were assessed using a Kruskal–Wallis test followed by Dunn’s post.

A heatmap of eDNA abundances was generated using ComplexHeatmap (v.2.25.2; Gu et al., 2016) from log10-transformed normalized abundance table, with a pseudocount of 1 × 10⁻⁶ added prior to transformation to accommodate zero values. All figures were generated using ggplot2 (v.3.5.1; Wickham, 2016), ggpubr (v.0.6.0; Kassambara, 2023), and patchwork (v.1.3.0; Pedersen, 2024). All R codes for figure generation are provided on GitHub (github.com/gogogogogul/). Results of statistical analyses are further documented in Supplementary Tables 10–14.

Apprill, A., McNally, S., Parsons, R., Weber, L., 2015. Minor revision to V4 region SSU rRNA 806R gene primer greatly increases detection of SAR11 bacterioplankton. Aquat. Microb. Ecol. 75, 129–137. https://doi.org/10.3354/ame01753

Arbizu, P.M., 2017. pairwiseAdonis: Pairwise Multilevel Comparison using Adonis.

Aroney, S.T.N., Newell, R.J.P., Nissen, J.N., Camargo, A.P., Tyson, G.W., Woodcroft, B.J., 2025. CoverM: read alignment statistics for metagenomics. Bioinformatics 41, btaf147. https://doi.org/10.1093/bioinformatics/btaf147

Bankevich, A., Nurk, S., Antipov, D., Gurevich, A.A., Dvorkin, M., Kulikov, A.S., Lesin, V.M., Nikolenko, S.I., Pham, S., Prjibelski, A.D., Pyshkin, A.V., Sirotkin, A.V., Vyahhi, N., Tesler, G., Alekseyev, M.A., Pevzner, P.A., 2012. SPAdes: a new genome assembly algorithm and its applications to single-cell sequencing. J. Comput. Biol. J. Comput. Mol. Cell Biol. 19, 455–477. https://doi.org/10.1089/cmb.2012.0021

Barnard, R.L., Osborne, C.A., Firestone, M.K., 2013. Responses of soil bacterial and fungal communities to extreme desiccation and rewetting. ISME J. 7, 2229–2241. https://doi.org/10.1038/ismej.2013.104

Bushnell, B., 2014. BBMap: A Fast, Accurate, Splice-Aware Aligner.

Callahan, B.J., McMurdie, P.J., Rosen, M.J., Han, A.W., Johnson, A.J.A., Holmes, S.P., 2016. DADA2: High-resolution sample inference from Illumina amplicon data. Nat. Methods 13, 581–583. https://doi.org/10.1038/nmeth.3869

Camargo, A.P., Roux, S., Schulz, F., Babinski, M., Xu, Y., Hu, B., Chain, P.S.G., Nayfach, S., Kyrpides, N.C., 2023. Identification of mobile genetic elements with geNomad. Nat. Biotechnol. 1–10. https://doi.org/10.1038/s41587-023-01953-y

Danecek, P., Bonfield, J.K., Liddle, J., Marshall, J., Ohan, V., Pollard, M.O., Whitwham, A., Keane, T., McCarthy, S.A., Davies, R.M., Li, H., 2021. Twelve years of SAMtools and BCFtools. GigaScience 10, giab008. https://doi.org/10.1093/gigascience/giab008

Emerson, J.B., Roux, S., Brum, J.R., Bolduc, B., Woodcroft, B.J., Jang, H.B., Singleton, C.M., Solden, L.M., Naas, A.E., Boyd, J.A., Hodgkins, S.B., Wilson, R.M., Trubl, G., Li, C., Frolking, S., Pope, P.B., Wrighton, K.C., Crill, P.M., Chanton, J.P., Saleska, S.R., Tyson, G.W., Rich, V.I., Sullivan, M.B., 2018. Host-linked soil viral ecology along a permafrost thaw gradient. Nat. Microbiol. 3, 870–880. https://doi.org/10.1038/s41564-018-0190-y

Eviner, V.T., Chapin, F.S., III, Vaughn, C.E., 2006. Seasonal Variations in Plant Species Effects on Soil N and P Dynamics. Ecology 87, 974–986. https://doi.org/10.1890/0012-9658(2006)87%5B974:SVIPSE%5D2.0.CO;2

Foley, M.M., Blazewicz, S.J., McFarlane, K.J., Greenlon, A., Hayer, M., Kimbrel, J.A., Koch, B.J., Monsaint-Queeney, V.L., Morrison, K., Morrissey, E., Hungate, B.A., Pett-Ridge, J., 2023. Active populations and growth of soil microorganisms are framed by mean annual precipitation in three California annual grasslands. Soil Biol. Biochem. 177, 108886. https://doi.org/10.1016/j.soilbio.2022.108886

Griffiths, R.I., Whiteley, A.S., O’Donnell, A.G., Bailey, M.J., 2000. Rapid Method for Coextraction of DNA and RNA from Natural Environments for Analysis of Ribosomal DNA- and rRNA-Based Microbial Community Composition. Appl. Environ. Microbiol. 66, 5488. https://doi.org/10.1128/aem.66.12.5488-5491.2000

Gu, Z., Eils, R., Schlesner, M., 2016. Complex heatmaps reveal patterns and correlations in multidimensional genomic data. Bioinformatics 32, 2847–2849. https://doi.org/10.1093/bioinformatics/btw313

Guo, J., Bolduc, B., Zayed, A.A., Varsani, A., Dominguez-Huerta, G., Delmont, T.O., Pratama, A.A., Gazitúa, M.C., Vik, D., Sullivan, M.B., Roux, S., 2021. VirSorter2: a multi-classifier, expert-guided approach to detect diverse DNA and RNA viruses. Microbiome 9, 37. https://doi.org/10.1186/s40168-020-00990-y

Hyatt, D., Chen, G.-L., LoCascio, P.F., Land, M.L., Larimer, F.W., Hauser, L.J., 2010. Prodigal: prokaryotic gene recognition and translation initiation site identification. BMC Bioinformatics 11, 119. https://doi.org/10.1186/1471-2105-11-119

John, S.G., Mendez, C.B., Deng, L., Poulos, B., Kauffman, A.K.M., Kern, S., Brum, J., Polz, M.F., Boyle, E.A., Sullivan, M.B., 2011. A simple and efficient method for concentration of ocean viruses by chemical flocculation. Environ. Microbiol. Rep. 3, 195–202. https://doi.org/10.1111/j.1758-2229.2010.00208.x

Kassambara, A., 2023. rstatix: Pipe-Friendly Framework for Basic Statistical Tests.

Langmead, B., Salzberg, S.L., 2012. Fast gapped-read alignment with Bowtie 2. Nat. Methods 9, 357–359. https://doi.org/10.1038/nmeth.1923

Li, H., Handsaker, B., Wysoker, A., Fennell, T., Ruan, J., Homer, N., Marth, G., Abecasis, G., Durbin, R., Subgroup, 1000 Genome Project Data Processing, 2009. The Sequence Alignment/Map format and SAMtools. Bioinformatics 25, 2078. https://doi.org/10.1093/bioinformatics/btp352

Muscatt, G., Hilton, S., Raguideau, S., Teakle, G., Lidbury, I.D.E.A., Wellington, E.M.H., Quince, C., Millard, A., Bending, G.D., Jameson, E., 2022. Crop management shapes the diversity and activity of DNA and RNA viruses in the rhizosphere. Microbiome 10, 181. https://doi.org/10.1186/s40168-022-01371-3

Nayfach, S., Camargo, A.P., Schulz, F., Eloe-Fadrosh, E., Roux, S., Kyrpides, N.C., 2021. CheckV assesses the quality and completeness of metagenome-assembled viral genomes. Nat. Biotechnol. 39, 578–585. https://doi.org/10.1038/s41587-020-00774-7

Nicolas, A.M., Sieradzki, E.T., Pett-Ridge, J., Banfield, J.F., Taga, M.E., Firestone, M.K., Blazewicz, S.J., 2023. A subset of viruses thrives following microbial resuscitation during rewetting of a seasonally dry California grassland soil. Nat. Commun. 14, 5835. https://doi.org/10.1038/s41467-023-40835-4

Nuccio, E.E., Anderson-Furgeson, J., Estera, K.Y., Pett-Ridge, J., de Valpine, P., Brodie, E.L., Firestone, M.K., 2016. Climate and edaphic controllers influence rhizosphere community assembly for a wild annual grass. Ecology 97, 1307–1318. https://doi.org/10.1890/15-0882.1

Nurk, S., Bankevich, A., Antipov, D., Gurevich, A.A., Korobeynikov, A., Lapidus, A., Prjibelski, A.D., Pyshkin, A., Sirotkin, A., Sirotkin, Y., Stepanauskas, R., Clingenpeel, S.R., Woyke, T., McLean, J.S., Lasken, R., Tesler, G., Alekseyev, M.A., Pevzner, P.A., 2013. Assembling single-cell genomes and mini-metagenomes from chimeric MDA products. J. Comput. Biol. J. Comput. Mol. Cell Biol. 20, 714–737. https://doi.org/10.1089/cmb.2013.0084

Oksanen, J., Simpson, G.L., Blanchet, F.G., Kindt, R., Legendre, P., Minchin, P.R., O’Hara, R.B., Solymos, P., Stevens, M.H.H., Szoecs, E., Wagner, H., Barbour, M., Bedward, M., Bolker, B., Borcard, D., Carvalho, G., Chirico, M., Caceres, M.D., Durand, S., Evangelista, H.B.A., FitzJohn, R., Friendly, M., Furneaux, B., Hannigan, G., Hill, M.O., Lahti, L., McGlinn, D., Ouellette, M.-H., Cunha, E.R., Smith, T., Stier, A., Braak, C.J.F.T., Weedon, J., Borman, T., 2025. vegan: Community Ecology Package.

Parada, A.E., Needham, D.M., Fuhrman, J.A., 2016. Every base matters: assessing small subunit rRNA primers for marine microbiomes with mock communities, time series and global field samples. Environ. Microbiol. 18, 1403–1414. https://doi.org/10.1111/1462-2920.13023

Paradis, E., Schliep, K., 2019. ape 5.0: an environment for modern phylogenetics and evolutionary analyses in R. Bioinformatics 35, 526–528. https://doi.org/10.1093/bioinformatics/bty633

Pedersen, T.L., 2024. patchwork: The Composer of Plots.

R Core Team, 2024. R: A Language and Environment for Statistical Computing. R Foundation for Statistical Computing, Vienna, Austria.

Roux, S., Adriaenssens, E.M., Dutilh, B.E., Koonin, E.V., Kropinski, A.M., Krupovic, M., Kuhn, J.H., Lavigne, R., Brister, J.R., Varsani, A., Amid, C., Aziz, R.K., Bordenstein, S.R., Bork, P., Breitbart, M., Cochrane, G.R., Daly, R.A., Desnues, C., Duhaime, M.B., Emerson, J.B., Enault, F., Fuhrman, J.A., Hingamp, P., Hugenholtz, P., Hurwitz, B.L., Ivanova, N.N., Labonté, J.M., Lee, K.-B., Malmstrom, R.R., Martinez-Garcia, M., Mizrachi, I.K., Ogata, H., Páez-Espino, D., Petit, M.-A., Putonti, C., Rattei, T., Reyes, A., Rodriguez-Valera, F., Rosario, K., Schriml, L., Schulz, F., Steward, G.F., Sullivan, M.B., Sunagawa, S., Suttle, C.A., Temperton, B., Tringe, S.G., Thurber, R.V., Webster, N.S., Whiteson, K.L., Wilhelm, S.W., Wommack, K.E., Woyke, T., Wrighton, K.C., Yilmaz, P., Yoshida, T., Young, M.J., Yutin, N., Allen, L.Z., Kyrpides, N.C., Eloe-Fadrosh, E.A., 2019. Minimum Information about an Uncultivated Virus Genome (MIUViG). Nat. Biotechnol. 37, 29–37. https://doi.org/10.1038/nbt.4306

Roux, S., Camargo, A.P., Coutinho, F.H., Dabdoub, S.M., Dutilh, B.E., Nayfach, S., Tritt, A., 2023. iPHoP: An integrated machine learning framework to maximize host prediction for metagenome-derived viruses of archaea and bacteria. PLOS Biol. 21, e3002083. https://doi.org/10.1371/journal.pbio.3002083

Roux, S., Emerson, J.B., Eloe-Fadrosh, E.A., Sullivan, M.B., 2017. Benchmarking viromics: an in silico evaluation of metagenome-enabled estimates of viral community composition and diversity. PeerJ 5, e3817. https://doi.org/10.7717/peerj.3817

Shaffer, M., Borton, M.A., McGivern, B.B., Zayed, A.A., La Rosa, S.L., Solden, L.M., Liu, P., Narrowe, A.B., Rodríguez-Ramos, J., Bolduc, B., Gazitúa, M.C., Daly, R.A., Smith, G.J., Vik, D.R., Pope, P.B., Sullivan, M.B., Roux, S., Wrighton, K.C., 2020. DRAM for distilling microbial metabolism to automate the curation of microbiome function. Nucleic Acids Res. 48, 8883–8900. https://doi.org/10.1093/nar/gkaa621

Sieradzki, E.T., Allen, G.M., Kimbrel, J.A., Nicol, G.W., Hazard, C., Nuccio, E., Blazewicz, S.J., Pett-Ridge, J., Trubl, G., 2025. Phosphate amendment drives bloom of RNA viruses after soil wet-up. Soil Biol. Biochem. 205, 109791. https://doi.org/10.1016/j.soilbio.2025.109791

Standard operating procedure for soil electrical conductivity, soil/water, 1:5, 2021.

Trubl, G., Roux, S., Borton, M.A., Varsani, A., Li, Y.-F., Sun, C.L., Jang, H.B., Woodcroft, B.J., Tyson, G.W., Wrighton, K.C., Saleska, S.R., Eloe-Fadrosh, E.A., Sullivan, M.B., Rich, V.I., 2025. Population ecology and biogeochemical implications of ssDNA and dsDNA viruses along a permafrost thaw gradient. Nat. Commun. 17, 375. https://doi.org/10.1038/s41467-025-67057-0

Trubl, G., Roux, S., Solonenko, N., Li, Y.-F., Bolduc, B., Rodríguez-Ramos, J., Eloe-Fadrosh, E.A., Rich, V.I., Sullivan, M.B., 2019. Towards optimized viral metagenomes for double-stranded and single-stranded DNA viruses from challenging soils. PeerJ 7, e7265. https://doi.org/10.7717/peerj.7265

Trubl, G., Solonenko, N., Chittick, L., Solonenko, S.A., Rich, V.I., Sullivan, M.B., 2016. Optimization of viral resuspension methods for carbon-rich soils along a permafrost thaw gradient. PeerJ 4, e1999. https://doi.org/10.7717/peerj.1999

Wickham, H., 2016. Introduction, in: Wickham, H. (Ed.), Ggplot2: Elegant Graphics for Data Analysis. Springer International Publishing, Cham, pp. 3–10. https://doi.org/10.1007/978-3-319-24277-4_1
